# Manganese acquisition is essential for virulence of *Enterococcus faecalis*

**DOI:** 10.1101/323832

**Authors:** C Colomer-Winter, AL Flores-Mireles, SP Baker, KL Frank, AJL Lynch, SJ Hultgren, T Kitten, JA Lemos

**Author notes:** Correspondence: 1395 Center Drive. PO Box 100424 Gainesville, FL, 32610, USA.

## Abstract

Manganese (Mn) is an essential micronutrient that is not readily available to pathogens during infection due to an active host defense mechanism known as nutritional immunity. To overcome this nutrient restriction, bacteria utilize high-affinity transporters that allow them to compete with host metal-binding proteins. Despite the established role of Mn in bacterial pathogenesis, little is known about the relevance of Mn in the pathophysiology of *E. faecalis*. Here, we identified and characterized the major Mn acquisition systems of *E. faecalis*. We discovered that the ABC-type permease EfaCBA and two Nramp-type transporters, named MntH1 and MntH2, work collectively to promote cell growth under Mn-restricted conditions. The simultaneous inactivation of EfaCBA, MntH1 and MntH2 (Δ*efa*Δ*mntH1*Δ*mntH2* strain) led to drastic reductions (>95%) in intracellular Mn content, severe growth defects in body fluids (serum and urine) *ex vivo*, significant loss of virulence in *Galleria mellonella*, and virtually complete loss of virulence in rabbit endocarditis and murine catheter-associated urinary tract infection (CAUTI) models. Despite the functional redundancy of *EfaCBA*, *MntH1* and *MntH2* under *in vitro* or *ex vivo* conditions and in the invertebrate model, dual inactivation of *efaCBA* and *mntH2* (Δ*efa*Δ*mntH2* strain) was sufficient to prompt maximal sensitivity to calprotectin, a Mn- and Zn-chelating host antimicrobial protein, and for the loss of virulence in mammalian models. Interestingly, EfaCBA appears to play a prominent role during systemic infection, whereas MntH2 was more important during CAUTI. The different roles of EfaCBA and MntH2 in these sites could be attributed, at least in part, to the differential expression of *efaA* and *mntH2* in cells isolated from hearts or from bladders. Collectively, this study demonstrates that Mn acquisition is essential for the pathogenesis of *E. faecalis* and validates Mn uptake systems as promising targets for the development of new antimicrobials.

## Author Summary

*Enterococcus faecalis* is a leading cause of hospital-acquired infections that are often difficult to treat due to their exceptional multidrug resistance. Manganese (Mn) is an essential micronutrient for bacterial pathogens during infection. To prevent infection, the host limits Mn bioavailability to invading bacteria in an active process known as nutritional immunity. To overcome this limitation, bacteria produce high-affinity Mn uptake systems to scavenge Mn from host tissues. Here, we identified the main Mn transporters of *E. faecalis* and show that, by working collectively, they are essential for growth of this opportunistic pathogen in Mn-restricted environments. Notably, the inability to acquire Mn during infection rendered *E. faecalis* virtually avirulent in different animal models, thereby revealing the essentiality of Mn acquisition to enterococcal pathogenesis. The results reported here highlight that bacterial Mn transport systems are promising targets for the development of novel antimicrobial therapies, which are expected to be particularly powerful to combat enterococcal infections.

## Introduction

While normal residents of the gastrointestinal (GI) tract of animals and humans, enterococci are also the third most frequent cause of hospital-acquired infections and a major threat to public health due to the alarming rise of multidrug-resistant isolates [1]. Enterococcal infections in humans are mainly caused by *Enterococcus faecalis* and *Enterococcus faecium*, with the great majority of infections (~70%) caused by *E. faecalis.* Collectively, enterococci rank as the third leading etiological agent in infective endocarditis (IE) [2], second in complicated urinary tract infections (UTI) [3], and one of the leading causes of device-associated infections and bacteremia [1]. Despite the recent introduction of new antibiotics active against both *E. faecalis* and *E. faecium* (i.e. daptomycin, linezolid, tigecycline), the indiscriminate use of antibiotics and the rise in elderly and severely ill populations susceptible to infection continues to contribute to a worldwide increase in enterococcal infections [4, 5]. The pathogenic potential of *E. faecalis*, and more generally, of all enterococci, has been largely attributed to their harsh and extremely durable nature, which includes intrinsic tolerance to commonly-used antibiotics (such as cephalosporins), chlorine, alcohol-based detergents, and an ability to survive extreme fluctuations in temperature, pH, oxygen tension, humidity and nutrient availability [1].

We recently showed that virulence-related phenotypes of a strain unable to synthesize the nutritional alarmone (p)ppGpp, known as (p)ppGpp^0^ strain, were directly linked to manganese (Mn) homeostasis, as important phenotypes of the (p)ppGpp^0^ strain could be reverted by addition of Mn to laboratory growth medium or serum [6]. Mn is an essential micronutrient for bacterial pathogens and hosts alike [7]. In lactic acid bacteria such as *E. faecalis*, Mn is the co-factor of enzymes of central metabolic pathways such as energy generation and DNA biosynthesis, and has been shown to play an important role in oxidative stress responses [8]. Similarly, vertebrates require Mn for a variety of cellular pathways such as lipid, protein, and carbohydrate metabolism [9, 10] and, depending on the tissue, Mn levels range from 0.3 to 2.9 μg per gram of tissue [11, 12]. While the lowest range concentration found in tissues is more than enough to promote bacterial growth, the mammalian host restricts the availability of essential metals such as Mn and iron (Fe) to invading pathogens by producing small molecules and proteins that tightly bind to metals, an active process termed nutritional immunity [13]. For example, Fe-binding proteins such as transferrin (TF) and lactoferrin (LF) are utilized by the host to chelate Fe in serum (TF) and mucose (LF), thereby restricting its bioavailability to invading pathogens [14]. The bioavailability of Mn and zinc (Zn) in the host is restricted, at least in part, by calprotectin, a Mn/Zn-sequestering protein of the S100 family that accounts for more than 40% of the total protein content of neutrophils [15]. In addition, macrophages express an Nramp (<u>N</u>atural <u>R</u>esistance-<u>A</u>ssociated <u>M</u>acrophage <u>P</u>rotein)-type Fe/Mn transporter that starves phagocytosed bacteria to promote clearance in the phagolysosome [7]. Notably, mice defective of calprotectin or the Nramp-type macrophage transporter are significantly more susceptible to bacterial infections [16, 17].

To overcome host-imposed Mn restriction, bacteria produce their own high-affinity Mn transporters. Presently, three classes of Mn transporters are known in bacteria: i) ABC-type permeases, ii) Nramp-type H^+^/Mn transporters, and iii) the less common P-type transporters [7]. Not surprisingly, Mn uptake systems have been identified as major virulence factors for several gram-negative and gram-positive pathogens [7, 8, 18]. For instance, in *Salmonella enterica* serovar Typhimurim, deletion of the ABC transporter *sitABCD* alone or in combination with the Nramp transporter *mntH* led to virulence attenuation in a murine systemic infection model [19]. Similarly, in *Staphylococcus aureus*, combined inactivation of the ABC-type transporter *mntABC* and the Nramp-type transporter *mntH* reduced staphylococcal virulence in murine skin abscess [20] and systemic infection models [17]. In streptococci, which are more closely related to the enterococci, deletion of the ABC-type Mn transporter attenuated virulence of *Streptococcus mutans, S. parasanguinis* and *S. sanguinis* in rat or rabbit models of IE as well as of *Streptococcus pyogenes* in a murine model of skin infection [21–25]. Remarkably, deletion of the ABC-type Mn transporter *psaBCA* rendered *Streptococcus pneumoniae* completely avirulent in at least three different animal models [26].

The significance of Mn homeostasis and specifically Mn acquisition systems to *E. faecalis* pathogenesis has not been explored in detail. Unlike the bacterial pathogens studied thus far that encode one or two high-affinity Mn transporters, *in silico* analysis indicates that the core genome of *E. faecalis* encodes three putative Mn transporters: the ABC-type transporter EfaCBA and two Nramp-type transporters designated MntH1 and MntH2 [27, 28]. Previous global transcriptional (microarray) analysis revealed that transcription of these genes, particularly *efaCBA* and *mntH2*, is strongly induced in blood or in urine *ex vivo* as well as in a murine peritonitis model [29–31]. In addition, transcription of *efaCBA, mntH2* and to a much lesser extent *mntH1* is induced in Mn-depleted laboratory medium while repressed in Mn-rich medium [6, 32–34]. Notably, an earlier study identified EfaA (the lipoprotein component of the EfaCBA complex) as a prominent antigen in enterococcal IE that can be used as an immunodiagnostic tool to discriminate *E. faecalis* from other IE-causing bacteria [35, 36]. Finally, virulence of a strain lacking *efaA* was slightly attenuated in the murine peritonitis model [36].

To determine the significance of Mn homeostasis in the pathophysiology of *E. faecalis*, we generated a panel of single (Δ*efa*, Δ*mntH1*, Δ*mntH2*), double (Δ*efa*Δ*mntH1*, Δ*efa*Δ*mntH2*, Δ*mntH1*Δ*mntH2*) and triple (Δ*efa*Δ*mntH1*Δ*mntH2*) deletion mutants in strain OG1RF and tested their ability to grow under metal-restricted conditions and to cause infection in three different animal models. We found that production of only one of the three Mn transporters is sufficient to support growth of *E. faecalis* in Mn-depleted media, a finding that is supported by the drastic reduction in cellular Mn levels in the triple Δ*efa*Δ*mntH1*Δ*mntH2* mutant but not in single or double mutant strains. Loss of *efaCBA* alone impaired growth in serum or in the presence of calprotectin *in vitro*. In most cases, inactivation of *mntH2* exacerbated the phenotypes of the Δ*efa* mutant, suggesting that EfaCBA and MntH2 are the primary high-affinity Mn transporters of *E. faecalis.* While only Δ*efa*Δ*mntH1*Δ*mntH2* showed virulence attenuation in the *Galleria mellonella* invertebrate model, both Δ*efal*Δ*mntH2* double and Δ*efa*Δ*mntH1*Δ*mntH2* triple mutant strains were virtually avirulent in the two mammalian models tested (i.e., rabbit IE and murine catheter-associated UTI). Collectively, this investigation highlights the essentiality of Mn acquisition to *E. faecalis* pathogenesis, suggesting that pathways associated with Mn homeostasis are promising targets for the development of new antimicrobials.

## Results

**EfaCBA, MntHl and MntH2 work in concert to promote growth of *E. faecalis* in Mn-restricted environments.** The annotations of EF2074-2076, EF1901 and EF1057 as *efaCBA, mntH1* and *mntH2*, respectively, have been inferred based on their similarities to previously characterized Mn transport systems and their Mn-dependent transcriptional profiles [32–34]. To uncover the role of these systems in metal uptake, single (Δ*efa*, Δ*mntH1*, Δ*mntH2*), double (Δ*efa*Δ*mntH1*, Δ*efamntH2*, Δ*mntH1* Δ*AmntH2*) and triple (Δ*efa*Δ*mntH1*Δ*AmntH2*) deletion strains were generated using a markerless system [37]. Brain Heart Infusion (BHI) broth has been previously reported to contain low levels of Mn [25]. Using inductively coupled plasma-optical emission spectrometry (ICP-OES), we confirmed this to also be the case for BHI prepared in our lab – Mn, Fe and Zn concentrations were ~ 60 ng ml^−1^ (Mn), ~ 220 ng ml^−1^ (Fe) and ~ 740 ng ml^−1^ (Zn) (Fig 1A). With those values in mind, all mutant strains were isolated on BHI plates supplemented with 150 μM MnSO_4_. Upon genetic confirmation that all gene deletions occurred as planned, we tested the capacity of all mutant strains to grow on unsupplemented BHI. While single and double mutant strains grew well on plain BHI plates, the triple Δ*efa*Δ*mntH1*Δ*mntH2* mutant could only grow on Mn-supplemented BHI (Fig 1B). To confirm this initial observation, growth of the parent OG1RF and mutant strains was monitored in chemically-defined FMC medium depleted of Mn (Mn < 5 ng ml^−1^) [6]. Growth of all mutant strains (singles, doubles and triple mutant strains) in complete FMC (Mn-replete) was indistinguishable from growth of the parent strain (Fig 1C, E). In contrast, in Mn-depleted FMC, the Δ*efa*Δ*mntH2* double mutant strain displayed an extended lag phase and slightly reduced final growth yields (Fig 1F). Most notably, both growth rate and final growth yields were severely impaired in the triple mutant strain (Fig 1F). Ectopic expression of any one of the three Mn transporters from a plasmid (pTG-*efa*, pTG-*mntH1* or pTG-*mntH2*) rescued the growth defect of the triple mutant on unsupplemented BHI agar and restored growth in Mn-depleted FMC (Fig 1G-H). Collectively, these results strongly support that EfaCBA, MntH1 and MntH2 are bona fide, functionally redundant, Mn transporters.

**Fig 1.**
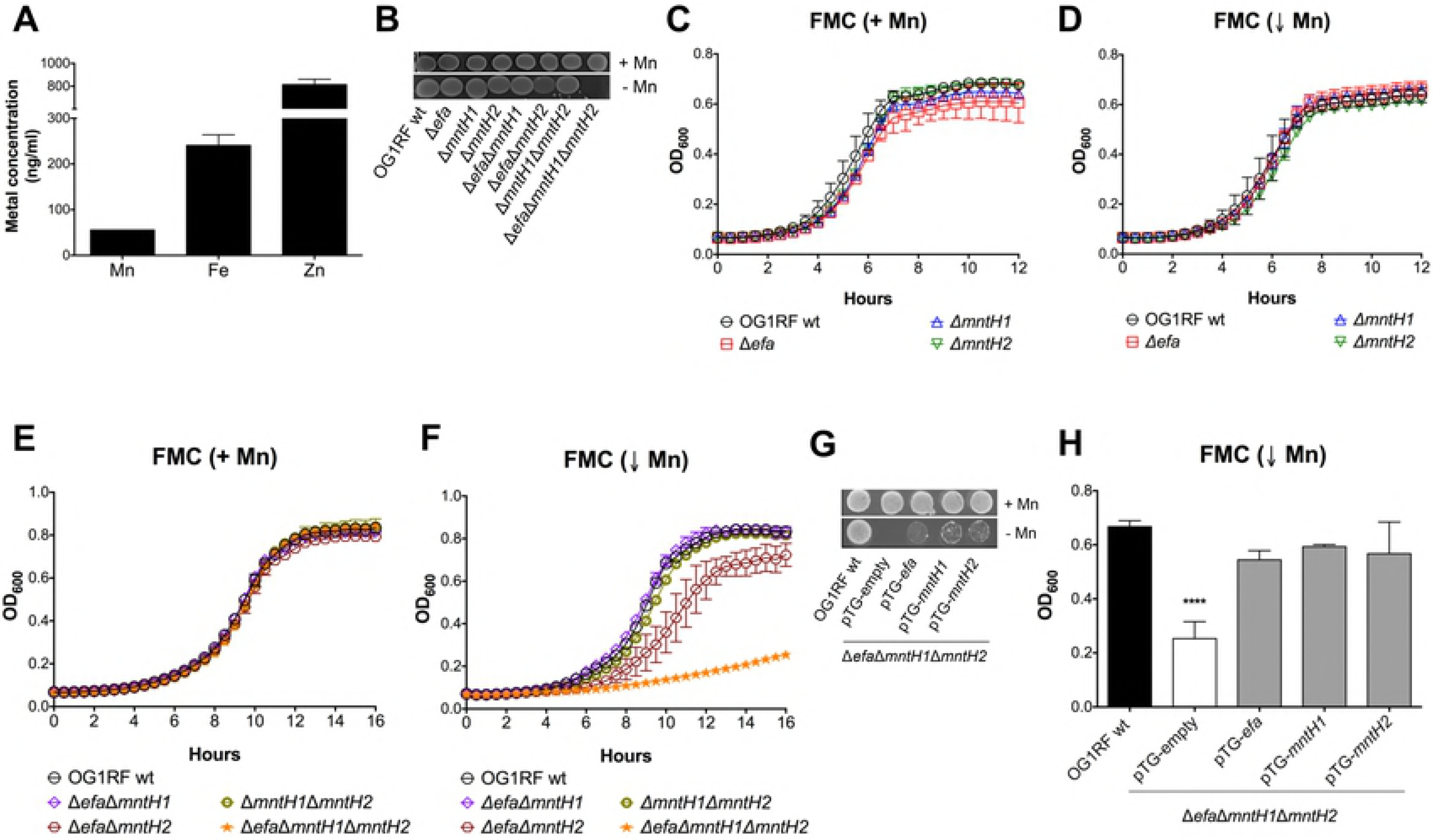
EfaCBA, MntH1 and MntH2 promote growth of *E. faecalis* in Mn-restricted environments. (A) Total Fe, Mn and Zn content of BHI medium determined by ICP-OES analysis. Error bars denote SD. (B) Growth of OG1RF (wild-type), Δ*efa*, Δ*mntH1*, Δ*mntH2*, Δ*efa*Δ*mntH1*, Δ*efa*Δ*mntH2*, Δ*mntH1*Δ*mntH2* and Δ*efa*Δ*mntH1*Δ*mntH2* strains on BHI plates with or without Mn supplementation. Overnight cultures were washed, diluted in PBS and 5 μl aliquots spotted on plates. Plates were incubated for 24 hours before being photographed. (C - F) Growth of OG1RF and its derivatives in Mn-replete (+ Mn, panels C, E) and Mn-depleted (↓ Mn, panels D, F) FMC. Cells were grown to OD_600_ ~ 0.2 in complete FMC and diluted 1:100 in either complete FMC or FMC depleted for Mn. Growth was monitored using a Bioscreen growth reader monitor. (G-H) Growth of OG1RF on BHI plates (G) or in Mn-depleted FMC broth (H) compared to the triple Δ*efa*Δ*mntH1*Δ*mntH2* mutant strain harboring either the empty plasmid (pTG-empty) or pTG-*efa*, pTG-*mntH1* or pTG-*mntH2*. Bars show optical density at 600 nm after 14 hours incubation. The graphs show the average and standard deviations of three independent experiments. Statistically significant differences were determined via ordinary one-way ANOVA with Dunnett’s multiple comparison post-test (\*\*\*\**p* ≤ 0.0001).

Next, we used ICP-OES to quantify cellular Mn content in the different Mn transport mutants. In agreement with the results shown in Figure 1, single deletions of *efaCBA, mntH1* or *mntH2* did not affect cellular Mn content (Fig 2A). However, combined deletion of *mntH2* with either *efaCBA* or *mntH1* (Δ*efa*Δ*mntH2* and Δ*mntH1*Δ*mntH2*) caused a significant reduction (~ 40%) in intracellular Mn pools (Fig 2B). Most notably, Mn pools were below detection limit in the triple mutant strain, thereby providing unequivocal evidence of the cooperative nature of EfaCBA, MntH1 and MntH2 in Mn acquisition. Taking into account that a balanced Mn/Fe ratio is essential for cellular homeostasis, and that bacterial systems homologous to EfaCBA have been shown to also transport Fe [7, 38], we again used ICP-OES to determine the cellular Fe content in our panel of mutant strains. While Fe pools were not affected in single and double mutants of Nramp-type transporters (Δ*mntH1*, Δ*mntH2* and Δ*mntH1*Δ*mntH2*), loss of EfaCBA, either alone or in combination with MntH1 and MntH2, led to ~ 40% reduction in cellular Fe content (Fig 2), indicating that, similar to streptococci, EfaCBA is a dual Mn/Fe transporter [7, 25].

**Fig 2.**
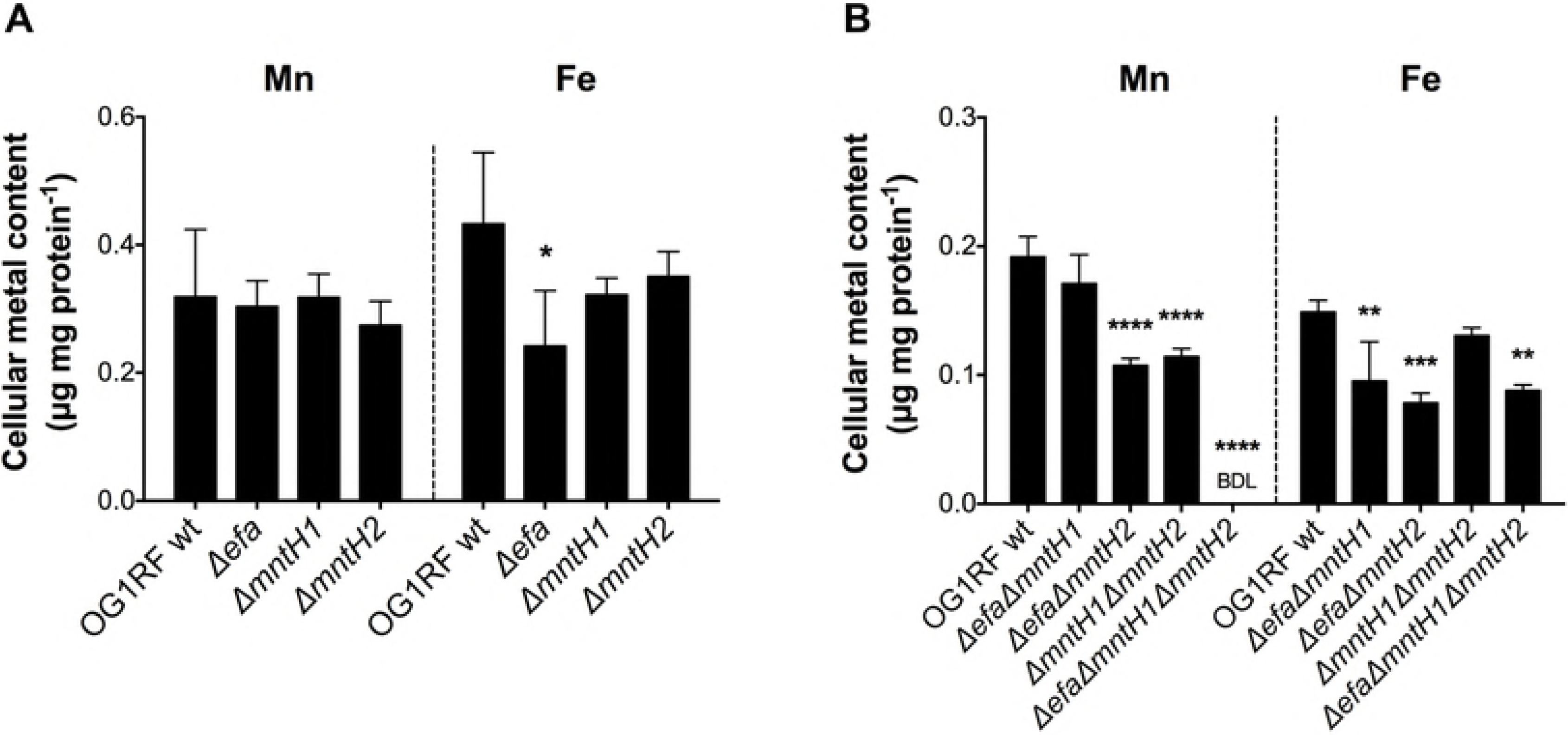
EfaCBA, MntH1 and MntH2 are the main Mn transporters of *E. faecalis.* Cellular Mn and Fe quantifications of *E. faecalis* OG1RF with (A) single Δ*efa*, Δ*mntH1*, Δ*mntH2*, or (B) double Δ*efa*Δ*mntH1*, Δ*efa*Δ*mntH2*, Δ*mntH1*Δ*mntH2* and triple Δ*efa*Δ*mntH1*Δ*mntH2* mutant strains. Strains were grown to mid-log phase (OD_600_ ~ 0.5) in BHI prior to analysis. The bar graphs show the average and standard deviations of three independent ICP-OES analyses. BDL: below detection limit. Statistically significant differences were determined via ordinary one-way ANOVA with Dunnett’s multiple comparison post-test (\**p* ≤ 0.05, ** *p* ≤ 0.01, \*\*\**p* ≤ 0.001, \*\*\*\**p* ≤ 0.0001).

**Virulence of the triple Δ*efa*Δ*mntH1*Δ*mntH2* strain is highly attenuated in the invertebrate *Galleria mellonella* model.** Metal sequestration is an ancient infection defense mechanism that spans various kingdoms of life [39]. Insects, such as *G. mellonella*, produce Fe-chelating proteins homologous to mammalian TF and ferritin and presumably use similar strategies to chelate Mn [40–42]. To probe the significance of Mn transport in *E. faecalis* virulence, we first used the *G. mellonella* larvae model of systemic infection. All single and double mutants killed *G. mellonella* at rates that were comparable to the parent OG1RF strain (Fig S1), with similar averages of larvae survival (~ 27%) 72 hours post-infection (Fig 3). However, virulence of the triple Δ*efa*Δ*mntH1*Δ*mntH2* mutant was dramatically impaired with ~ 87% larvae survival after 72 hours (Fig 3, Fig S1).

**Fig 3.**
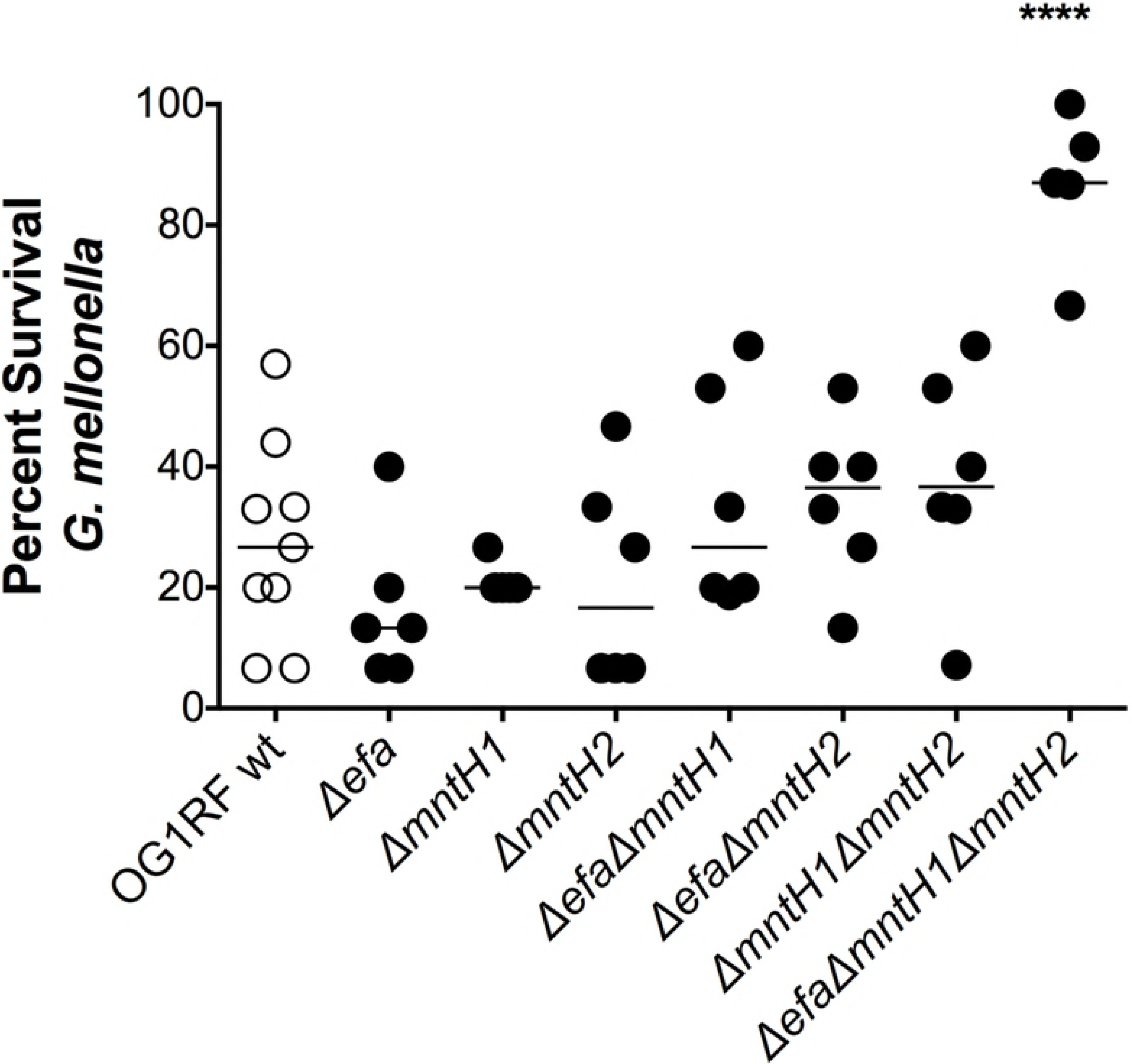
Virulence of the triple Δ*efaCBA*Δ*mntH1*Δ*mntH2* strain is attenuated in the *G. mellonella* model. Percent survival of *G. mellonella* 72 hours post-infection. Each symbol corresponds to a group of 15 larvae injected with ~ 5 × 10^5^ CFU of *E. faecalis.* Mean value is shown as a horizontal line. Final 72 hours survival of larvae infected with different strains was compared via ordinary one-way ANOVA with Dunnett’s multiple comparison post-test. (\*\*\*\**p* ≤ 0.0001).

**The EfaCBA system, and to a lesser extent MntH2, are required for calprotectin tolerance.** The Mn- and Zn-binding calprotectin is a major protein used by vertebrates to restrict the availability of these two transition metals to microbes during infection [15]. Normally found in circulating blood and tissues at relatively low levels, calprotectin rapidly accumulates to concentrations of up to 1mg ml^−1^ in response to inflammation and infection [15], thereby depleting Zn and Mn from the infected site. Of note, previous studies indicated that the antimicrobial activity of calprotectin against *E. faecalis* is, at least in part, due to Mn sequestration [15]. Here, we assessed the ability of the Mn transport mutants to grow in the presence of a sub-inhibitory concentration of purified calprotectin over a 24 hours period. Loss of *mntH1* or *mntH2* did not affect (Δ*mntH1*) or modestly affected (Δ*mntH2*) growth of *E. faecalis* in the presence of calprotectin (Fig 4). On the other hand, growth of any one of the *efaCBA* mutants (Δ*efa*, Δ*efa*Δ*mntH1*, Δ*efa*Δ*mntH2* and Δ*efa*Δ*mntH1*Δ*mntH2*) was strongly impaired (>50% growth inhibition) in the presence of calprotectin. As expected, double inactivation of *efaCBA* and *mntH2* (Δ*efa*Δ*mntH2*) rendered *E. faecalis* even more susceptible to the inhibitory effects of calprotectin. These results revealed that EfaCBA, closely followed by MntH2, is the primary Mn transporter used by *E. faecalis* to overcome the inhibitory effects of calprotectin.

**Fig 4.**
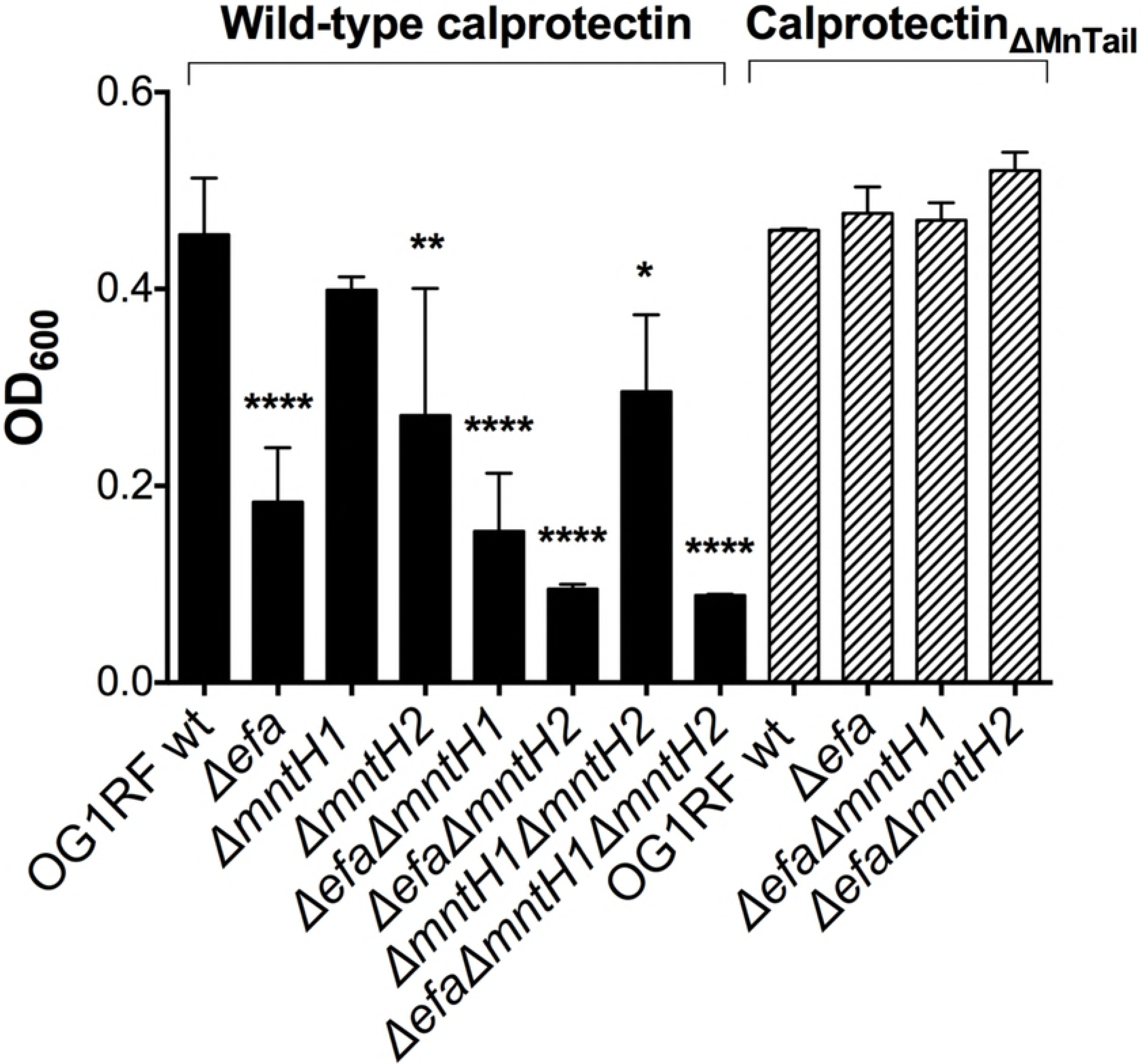
EfaCBA and MntH2 are required for *E. faecalis* tolerance to calprotectin. Growth of OG1RF and its derivatives in the presence of purified calprotectin (wild-type) or a calprotectin variant unable to chelate Mn (Calprotectin_ΔMn Tail_). Overnight cultures were diluted 1:50 in BHI and incubated at 37°C for 1 hour prior to diluting 1:100 into CP medium and incubating the strains with 120 μg ml^−1^ of purified calprotectin. Growth was monitored using a Bioscreen growth reader monitor for up to 24 hours. The bar graphs show the average and standard deviations of three independent cultures. Growth differences in the presence of wild-type calprotectin were analyzed via a two-way ANOVA with Tukey’s comparison post-test. (\**p* ≤ 0.05, \*\**p* ≤ 0.01, \*\*\*\**p* ≤ 0.0001).

A member of the S100 family, calprotectin forms a S100A8/S100A9 heterodimer with two distinct metal-binding sites: a high-affinity Zn-binding site and a dual high-affinity Zn/Mn-binding site. To confirm that the increased sensitivity of Δ*efa* strains to calprotectin was due to Mn sequestration and not to Zn sequestration, we tested the sensitivity of wild-type and selected mutant strains against a calprotectin variant (ΔMn Tail calprotectin) that retains the ability to chelate Zn but is unable to chelate Mn due to amino acid substitutions of key histidine residues (His103 and His105) in the C-terminal tail of calprotectin [15]. The inability to chelate Mn by the calprotectin ΔMn Tail variant restored final growth yields of all mutant strains tested to wild-type levels (Fig 4), indicating that EfaCBA, and to a lesser extent MntH2, promote tolerance to calprotectin in a Mn-dependent manner.

**EfaCBA is central for growth and survival in serum.** Next, we monitored growth and survival of the Mn transport mutants in pooled human serum at 37°C – serum was used instead of whole blood to avoid free metal contamination released by lysing erythrocytes. While inactivation of *mntH1* and *mntH2*, alone or in combination, did not affect the ability of *E. faecalis* to grow and survive in serum, loss of *efaCBA* led to a significant decrease in survival after 48 hours incubation when compared to the parent strain (~1.3 log Δ*efa*, ~1.5 log Δ*efa*Δ*mntH1*, ~2.3 log Δ*efa*Δ*mntH2*, ~1.8 log, Δ*efa*Δ*mntH1*Δ*mntH2*) (Fig 5 A-B). To determine if the survival defect of Δ*efa* strains was due to an inability to scavenge Mn or Fe from the environment, we supplemented serum with 1 mM MnSO_4_ or 1 mM FeSO_4_. The addition of either Mn or Fe restored serum growth and survival of Δ*efa* strains to wild-type OG1RF levels (Fig 5C). Moreover, Mn or Fe supplementation increased final growth yields of all strains, including the parent strain, confirming that both metals are growth limiting factors in the human serum. Collectively, these results indicate that the EfaCBA system plays a primary role in promoting *E. faecalis* serum survival, likely because of its dual capacity to function as a Mn and Fe transporter.

**Fig 5.**
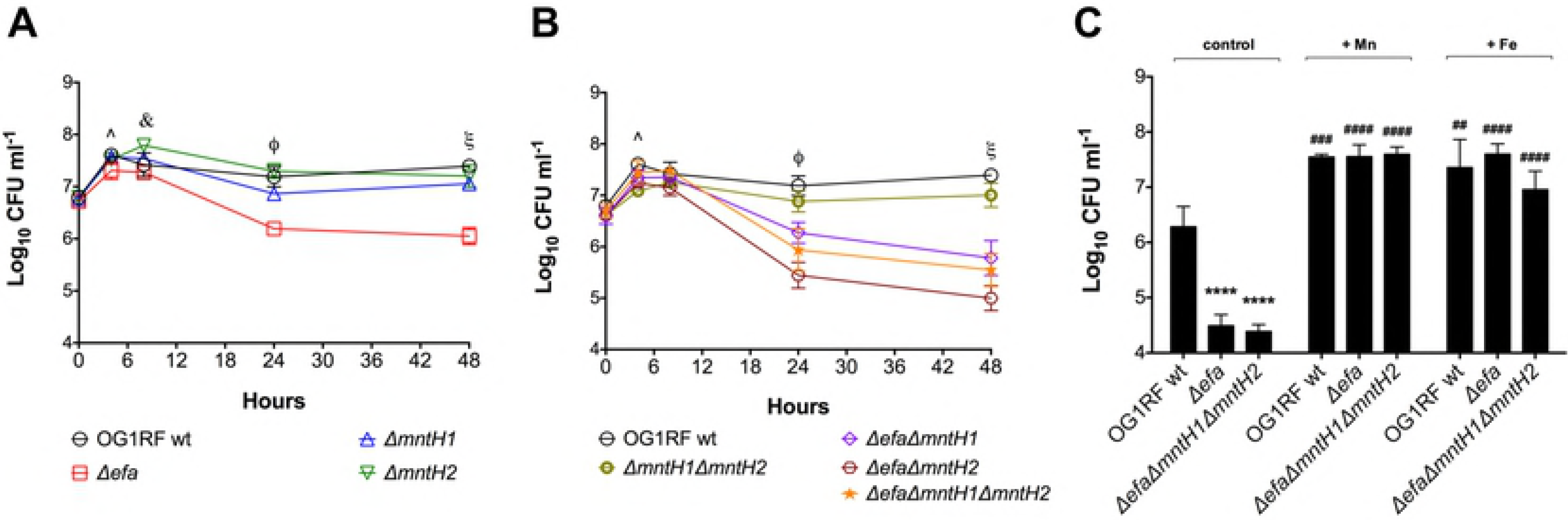
Growth and survival profile of *E. faecalis* in human serum. Growth and survival of *E. faecalis* OG1RF with (A) single Δ*efa*, Δ*mntH1*, Δ*mntH2* or (B) double Δ*efa*Δ*mntH1*, Δ*efa*Δ*mntH2*, Δ*mntH1*Δ*mntH2*, and triple Δ*efa*Δ*mntH1*Δ*mntH2* mutant strains in pooled human serum. (C) Serum was supplemented with 1 mM FeSO_4_ or 1 mM MnSO_4_, and survival of OG1RF, Δ*efa* and Δ*efa*Δ*mntH1*Δ*mntH2* strains was recorded after 48 hours incubation. Control corresponds to serum without metal supplementation. Aliquots at selected time points were serially diluted and plated on BHI + Mn plates for CFU enumeration. The graphs show the average log_10_-transformed CFU mean and standard deviations of at least three independent experiments. In (A-B), mutant strains were compared against wild-type OG1RF by a two-way ANOVA with Dunnett’s multiple comparison test. (\**p* ≤ 0.05, \*\**p* ≤ 0.01,\*\*\*\**p* ≤ 0.0001). Significant differences at (^) 4 hours of incubation: *Δ*efa*, *Δ*efa*Δ*mntH2*, **Δ*mntHl*Δ*mntH2*; at (&) 8 hours of incubation: **Δ*mntH2*; at (ϕ) 24 hours of incubation: ****Δ*efa*, **Δ*mntHl* ****Δ*efa*Δ*mntHl*, ****Δ*efa*Δ*mntH2*, ****Δ*efa*Δ*mntHl*Δ*mntH2*; at (ξ) 48 hours of incubation: ****Δ*efa*, **Δ*mntHl*, ****Δ*efa*Δ*mntHl*, ****Δ*efa*Δ*mntH2*, *Δ*mntHl* Δ*mntH2*, **** Δ*efa*.Δ*mntHl* Δ*mntH2*. In (C), comparison of survival defects among strains and the effect of Fe or Mn supplementation on the same strain (#) versus untreated, or between parent and mutant strains (*) under the same conditions were assessed via two-way ANOVA with Tukey’s post-test on log_10_-transformed CFU values. (##*p* ≤ 0.01, ###*p* ≤ 0.001, **** or ####*p* ≤ 0.0001).

**EfaCBA and MntH2 contribute to virulence in a rabbit model of IE.** Patients with enterococcal IE are known to generate specific antibodies against EfaA, the substrate-binding lipoprotein component of the EfaCBA system [32, 35]. While an *efaA* single mutant was previously shown to be slightly attenuated in a mouse peritonitis model [36], the contribution of EfaCBA, or the other Mn transport systems MntH1 and MntH2, in enterococcal IE has not been explored. Here, we used a catheterized rabbit IE model [43] to determine the ability of selected Mn transport mutants to colonize a previously-formed sterile heart vegetation and, then, systemically spread to different organs by using spleen homogenates as a readout. In the first experimental set, the parent OG1RF strain was co-inoculated systemically (via ear vein injection) with single Δ*efa* and triple Δ*efa*Δ*mntH1*Δ*mntH2* strains. Forty-eight hours post-infection, an average of 7.5 (+/− 0.7 SD) and 4.1 (+/− 0.9 SD) log_10_ CFU *E. faecalis* were recovered from hearts and spleens, respectively, confirming that *E. faecalis* can efficiently colonize the injured heart endothelium and spread systemically in this model (data not shown). To avoid any bias or inconsistencies that could be generated by antibiotic cassette markers, PCR was used to screen *E. faecalis* colonies (50 to 100 per animal and organ) to distinguish between the three strains based on the different amplification products obtained using *efaCBA* and *mntH2* flanking primers. Of the total colonies screened from heart vegetations, ~ 55% corresponded to the parent OG1RF and ~ 45% to the Δ*efa* single mutant strain (Fig 6A). While the average recovery rates of OG1RF and Δ*efa* were not statistically significant (*p*>0.05), there was a great variation from animal to animal, with the parent OG1RF corresponding to the large majority (>90%) of colonies recovered in two animals and Δ*efa* predominating at ~ 80% in the remaining three animals (Fig 6A). Similar trends were observed in the corresponding spleens of the individual animals (~ 49% OG1RF and ~ 51% Δ*efa*, *p*>0.05) (Fig 6B), indicating that systemic dissemination is likely a direct consequence of bacterial seeding from the heart vegetation. Most importantly, none of the screened colonies in this experiment corresponded to the triple mutant strain in either heart vegetations or spleens (Fig 6A-B).

**Fig 6.**
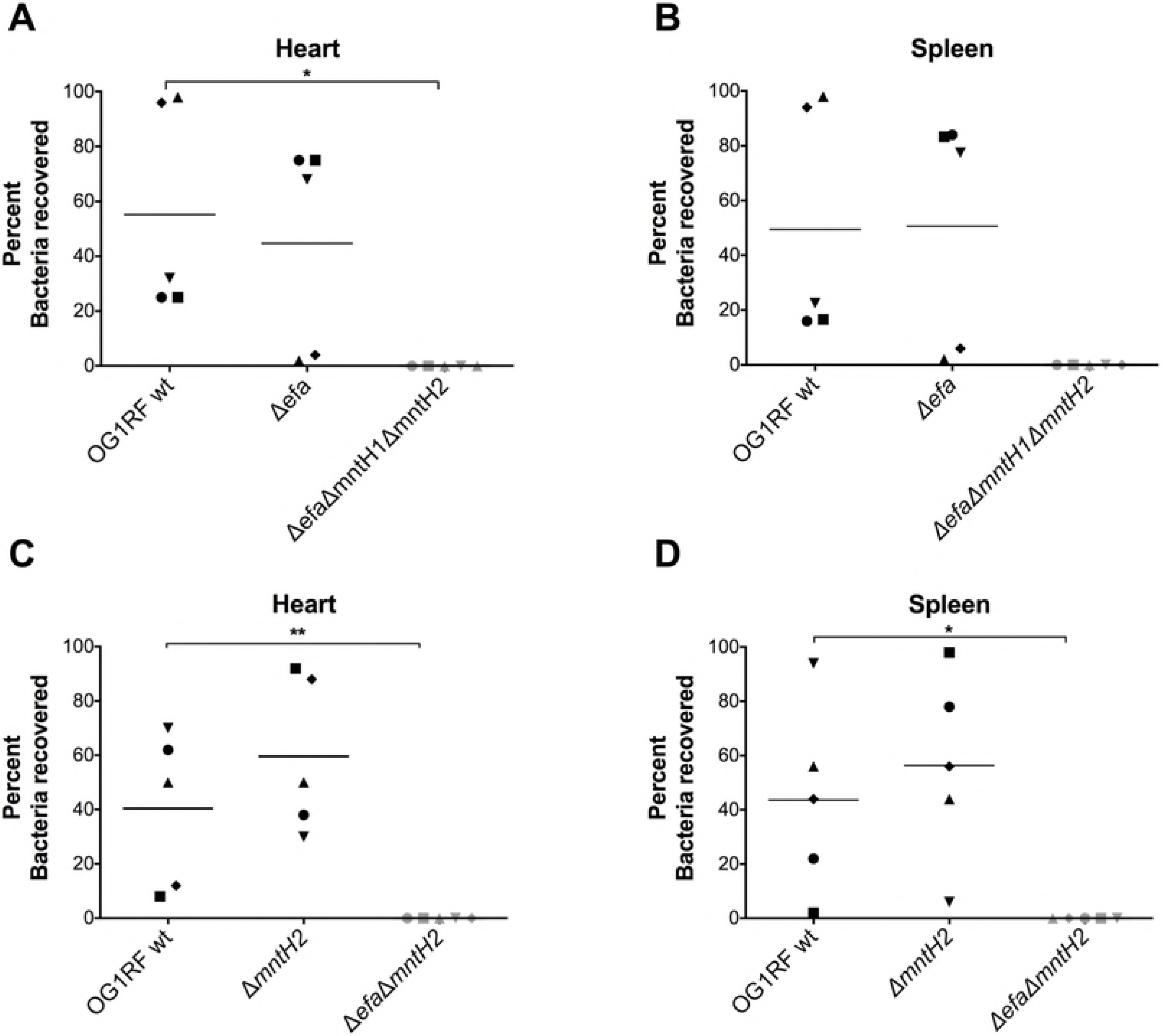
EfaCBA and MntH2 contribute to virulence in a rabbit model of IE. OG1RF and its derivatives (A-B) Δ*efa* and Δ*efa*Δ*mntH1*Δ*mntH2* or (C-D) Δ*mntH2* and Δ*efa*Δ*mntH2* were co-inoculated (1:1:1 ratio) into the ear vein of rabbits 24 hours after catheter implantation. After 48 hours, animals were euthanized and bacterial burdens determined in (A, C) hearts and (B, D) spleens. Graphs show the percent of each strain recovered from these sites; each symbol represents an individual rabbit. Symbols in grey indicate that this particular strain was not recovered from the corresponding animal. The mean percent recovery is shown as a horizontal line. Statistically significant differences in the percent of individual strains recovered from the rabbit IE experimental model were determined using an ordinary one-way ANOVA with Tukey’s comparison post-test. (\**p* ≤ 0.05, ** *p* ≤ 0.01).

Based on the strong growth defect of the double Δ*efa*Δ*mntH2* in the presence of calprotectin and in serum (Fig 4 and 5), we next tested the ability of the parent OG1RF, Δ*mntH2* and Δ*efa*Δ*mntH2* strains to colonize the heart vegetations in a triple co-infection experiment. In this second set of experiments, the average numbers of OG1RF and Δ*mntH2* recovered from heart vegetations and spleens were not significantly different (~ 40% OG1RF, ~ 60% Δ*mntH2*, *p*>0.05) (Fig 6C-D). Interestingly, the double Δ*efa*Δ*mntH2* mutant phenocopied the triple mutant strain, as none of the colonies screened from each animal corresponded to the Δ*efa*Δ*mntH2* strain. Collectively, these results strongly support that EfaCBA and MntH2 are the primary Mn transporters of *E. faecalis* during infection and further underscore the functional redundancy shared by these two Mn transporters *in vivo*.

**EfaCBA and MntH2 can individually support growth of *E. faecalis* in urine.** Previous studies have shown that transcription of *efaCBA* and *mntH2* is induced when *E. faecalis* is grown in urine, indicating that Mn is also a growth-limiting nutrient in the bladder environment [30, 44]. We used ICP-OES to determine the Mn content in pooled human urine obtained from healthy donors and found this batch to be low in Mn (~ 40 ng ml^−1^), which is within the normal clinical range described elsewhere [10]. Copper (Cu) (~ 10 ng ml^−1^) and Fe (~ 40 ng ml^−1^) levels were also low in pooled urine, whereas Zn levels (~ 600 ng ml^−1^) were comparatively much higher. Nevertheless, it should be noted that these values represent total metal concentrations present in urine not taking into account their bioavailability. Next, we tested the ability of the mutant strains to grow in human urine supplemented with bovine serum albumin (BSA) at 37°C – BSA was used to supplement urine to mimic protein infiltration during catheter-associated urinary tract infections (CAUTI), a condition shown to promote growth of *E. faecalis* in the bladder environment [45, 46]. We found that both the double Δ*efa*Δ*mntH2* and triple Δ*efa*Δ*mntH1*Δ*mntH2* mutant strains displayed a significant growth defect after 24 hours of incubation (Fig 7A). While the parent, single and double mutant strains entered stationary phase and maintained the same growth yields for up to 48 hours, the Δ*efa*Δ*mntH1*Δ*mntH2* triple mutant also displayed a survival defect after 48 hours (~ 1 log_10_ reduction in CFU ml^−1^, *p* < 0.05) when compared to the other strains (data not shown). Importantly, Mn supplementation fully restored growth of the triple mutant strain in urine, while addition of Fe only partially rescued the defective phenotype (Fig 7B).

**Fig 7.**
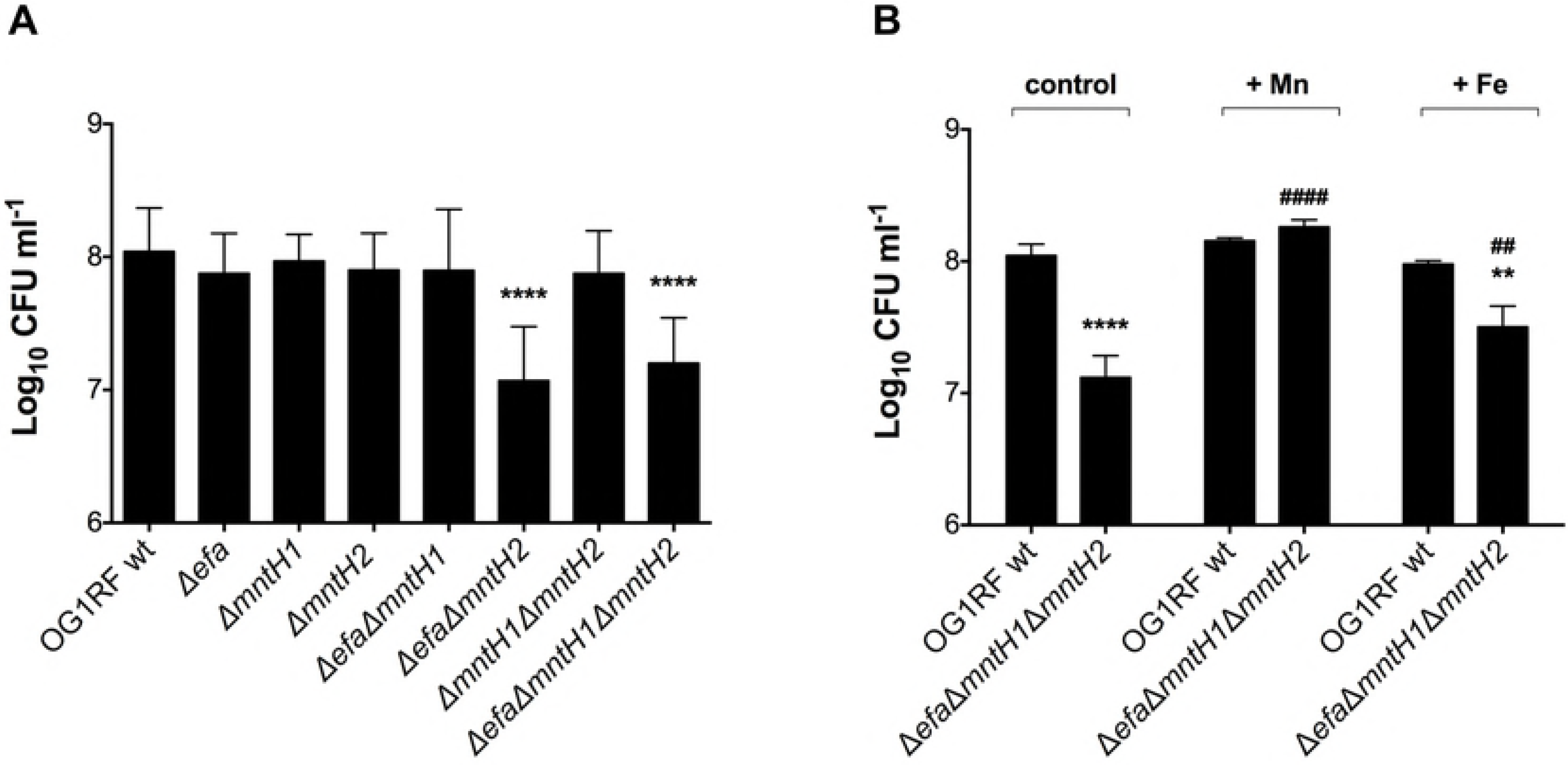
Growth and survival of *E. faecalis* in human urine. (A) Growth of OG1RF and derivatives in human urine. Overnight cultures were diluted 1:1000 into pooled urine supplemented with 20 mg ml^−1^ BSA and incubated for 24 hours at 37°C. (B) Human urine was supplemented with 1 mM FeSO_4_ or 1 mM MnSO_4_ and growth/survival of wild-type and Δ*efa*Δ*mntH1*Δ*mntH2* strains recorded after 48 hours incubation. Control corresponds to urine without metal supplementation. Aliquots at selected time points were serially diluted and plated on BHI + Mn plates for CFU enumeration. The graphs show the average and standard deviations of three independent experiments. Mutant strains were compared against wild-type OG1RF using one-way ANOVA followed by Sidak’s multiple comparison test. Comparison of the effect of Fe or Mn supplementation on the same strain (#) versus untreated, or between parent and mutant strain (*) under the same conditions, were assessed via two-way ANOVA with Tukey’s post-test. (** and ##*p* ≤ 0.01, **** and ####*p* ≤ 0.0001).

**MntH2, and to a lesser extent EfaCBA, contribute to *E. faecalis* virulence in a murine CAUTI model.** To determine whether Mn acquisition is also relevant to enterococcal UTI, we tested the parent OG1RF and each individual mutant strain in a murine CAUTI model [47]. Twenty-four hours post-infection, the OG1RF, Δ*efa* and Δ*mntH1* strains were recovered in similar numbers from the bladders of infected animals, whereas the Δ*mntH2* strain had a ~ 1.5 log_10_ CFU reduction (*p*<0.05) in total bacteria recovered (Fig 8A). Inactivation of *mntH1* did not exacerbate the phenotype of the *mntH2* single gene inactivation (Δ*mntH1* Δ*mntH2*) but simultaneous inactivation of *efaCBA* and *mntH1*(Δ*efa*Δ*mntH1*) resulted in ~ 1 log_10_ CFU reduction in bacteria recovered from the bladder (*p*<0.05). In agreement with the mounting evidence supporting that EfaCBA and MntH2 are the primary systems for Mn acquisition, combined deletion of *efaCBA* and *mntH2* (Δ*efa*Δ*mntH2*) or all three Mn transporters (Δ*efa*Δ*mntH1*Δ*mntH2*) further reduced bacterial loads recovered from the bladder to below or near the detection limit (Fig 8A). The exact same trends found in the bladder were observed for bacteria recovered from biofilms formed on the surface of the implanted catheters, i.e. significantly lower bacterial burden for strains lacking *mntH2* and very little to no bacterial cells recovered from strains lacking both *efaCBA* and *mntH2* (Fig 8B). Moreover, while the parent OG1RF was able to ascend to the kidneys in 55% of the mice and disseminate to spleen and heart in ~ 30% of the animals infected, the Mn transport mutants were rarely recovered from kidneys and were almost never isolated from more distant organs such as spleen and heart (Fig S2).

**Fig 8.**
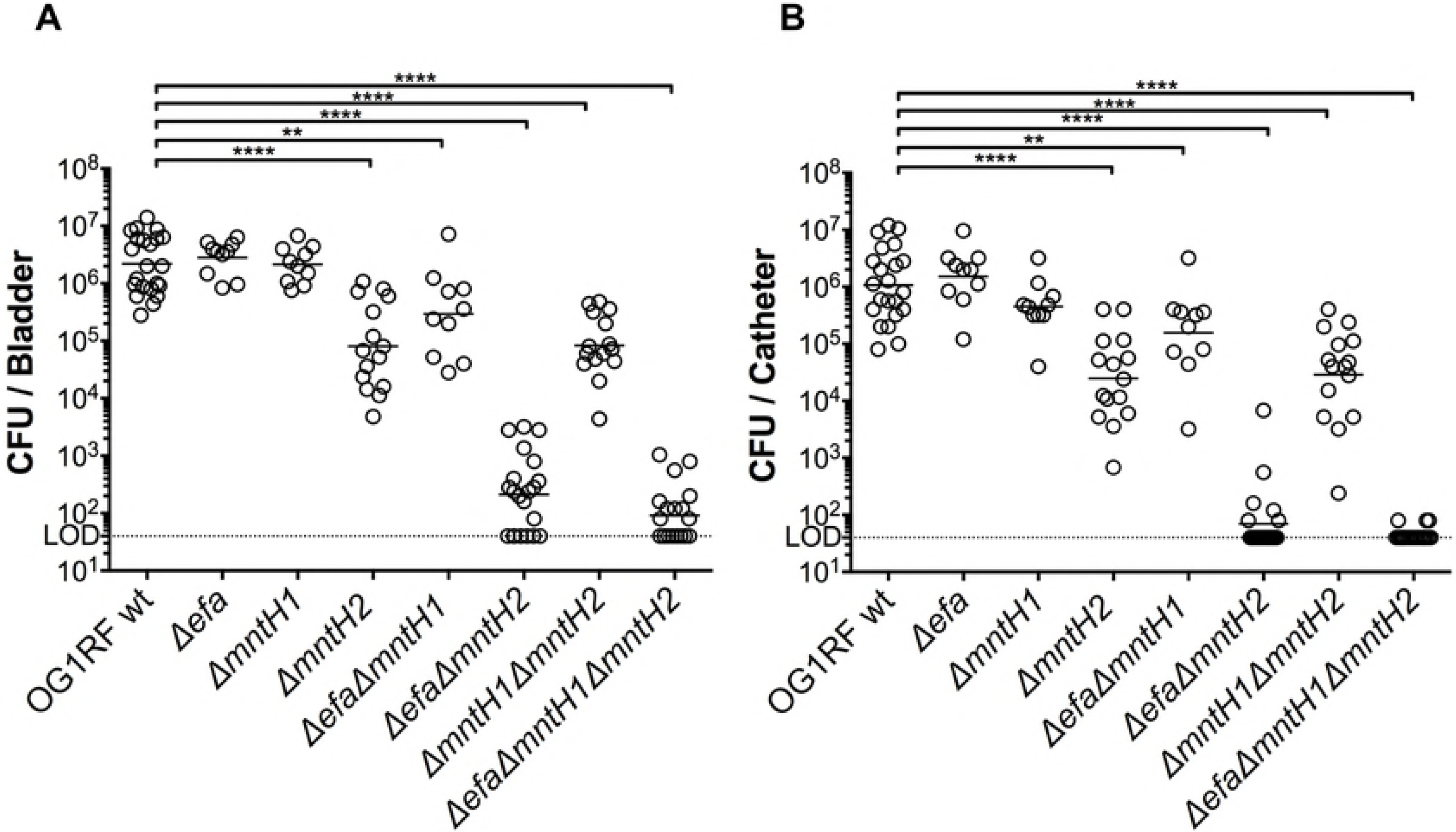
MntH2 and EfaCBA promote virulence of *E. faecalis* in a murine CAUTI model. OG1RF and its derivatives were inoculated into the bladder of mice immediately after catheter implantation (n = 10 to 22). After 24 hours, animals were euthanized, and bacterial burdens in (A) bladders and (B) catheters determined. Graphs show total CFU recovered from these sites, and each symbol represents an individual mouse. Symbols on the dashed line indicate that recovery was below the limit of detection (LOD, 40 CFU). Data were pooled from at least two independent experiments, and the median value is shown as a horizontal line. Two-tailed Mann-Whitney *U* tests were performed to determine significance (\*\**p* < 0.005, \*\*\*\**p* < 0.0001).

**Mn acquisition promotes biofilm formation in urine.** Biofilm formation of *E. faecalis* on urinary catheters is a critical step in UTI and is primarily mediated by the enterococcal surface adhesin EbpA [46, 48]. Notably, the metal ion-dependent adhesion site (MIDAS) of EbpA is required for biofilm formation in urine and for virulence in the experimental CAUTI model [46, 49]. While the identity of the metal bound to the tip of EbpA remains elusive, we wondered if the colonization defect of strains lacking one or more Mn transporters could relate to impaired biofilm formation and/or defects in EbpA production. First, we tested the ability of the parent and mutant strains to form biofilms on the surfaces of tissue culture plate wells and plastic catheters that were pre-coated with fibrinogen to promote biofilm formation in an EbpA-dependent manner [46]. After 24 hours of incubation in urine supplemented with BSA, the single Δ*efa*, double Δ*efa*Δ*mntH2* and triple Δ*efa*Δ*mntH1*Δ*mntH2* mutant strains formed significantly less biofilm as measured by either crystal violet staining of tissue culture plates (Fig 9A) or catheter immunostaining (Fig 9B). In agreement with these findings, the double Δ*efa*Δ*mntH2* and triple Δ*efa*Δ*mntH1*Δ*mntH2* mutant strains displayed a ~ 1 log_10_ CFU reduction in catheter-associated bacteria when compared to the parent strain (Fig 9C). While compatible with the slight reduction in biofilm formation (Fig 9A-B), the ~ 0.5 log_10_ CFU reduction observed with the Δ*efa* single mutant on the catheter surface was not statistically significant (Fig 9C). Finally, we used ELISA to quantify the surface expression levels of EbpA in the different strains grown under the same conditions used in the biofilm assay, i.e. urine supplemented with BSA. Despite defects in biofilm formation of Δ*efa*, Δ*efa*Δ*mntH2* and Δ*efa*Δ*mntH1*Δ*mntH2* strains on fibrinogen-coated surfaces, there were no apparent differences in EbpA levels between parent and mutant strains (Fig 9D), suggesting that the defective biofilm phenotype is not EbpA-dependent.

**Fig 9.**
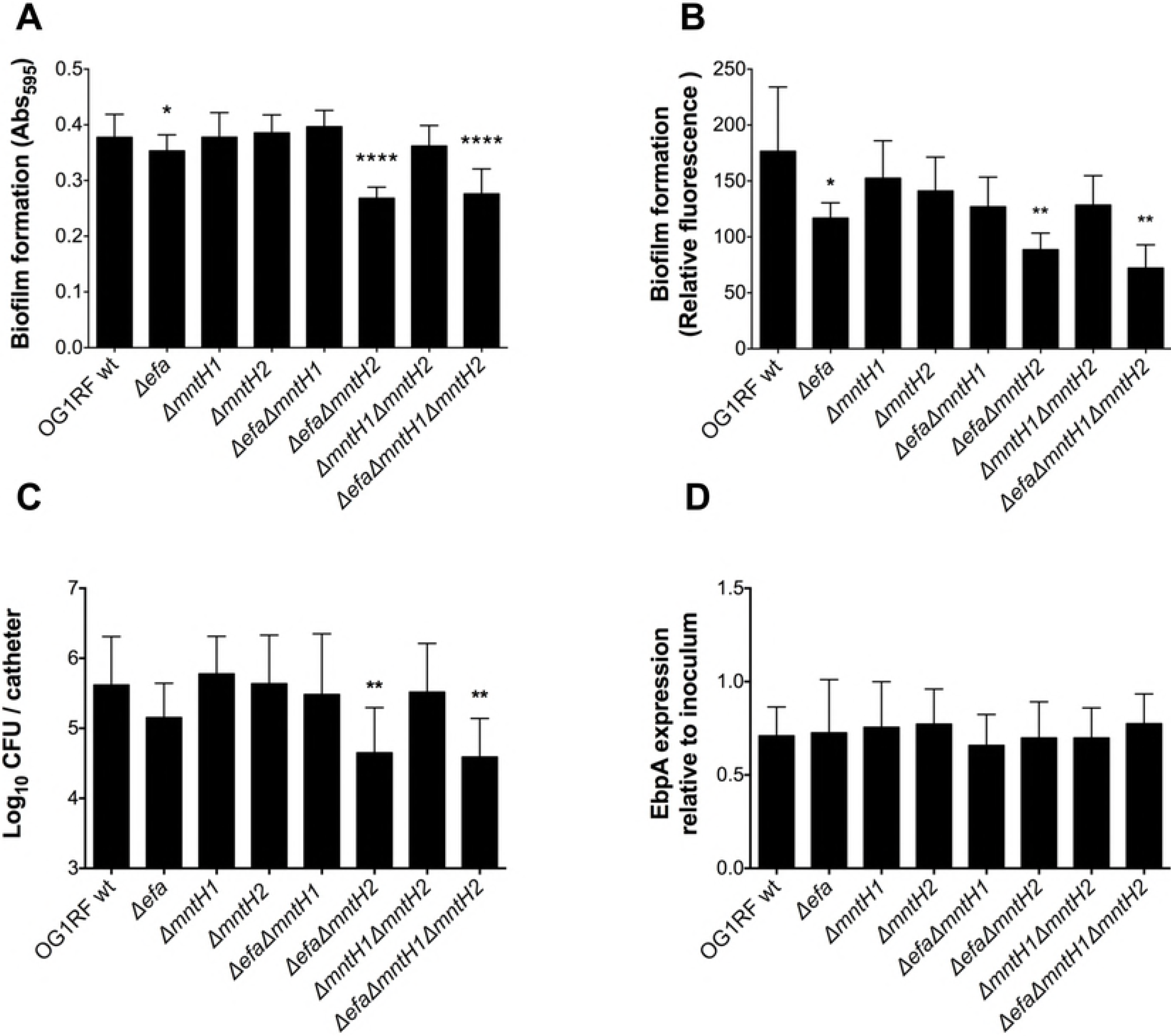
High-affinity Mn acquisition promotes biofilm formation in urine. (A-B) Fibrinogen-coated (A) 96-well polystyrene plates or (B) silicone catheters were incubated with *E. faecalis* strains for 24 hours in human urine supplemented with 20 mg ml^−1^ BSA. (A) Plates were stained with 0.5% crystal violet, which was dissolved with 33% acetic acid and absorbance at 595 nm was measured. (B) Fixed catheters were incubated with rabbit anti-*Streptococcus* group D antigen and Odyssey secondary antibody. Biofilm formation was quantified by infrared scanning. (C) Catheters were vortexed and sonicated to retrieve biofilm-associated bacteria and plated for cell enumeration. (D) EbpA expression in urine. Strains were grown in urine + BSA overnight prior to quantification of EbpA by ELISA. EbpA surface expression was detected using mouse anti-EbpA^Full^ and HRP-conjugated goat anti-rabbit antisera, and absorbance was determined at 450 nm. EbpA expression titers were normalized against the bacterial titers. Experiments were performed independently in triplicate per condition and per experiment. Biofilm quantification assays and EbpA expression were analyzed by a two-tailed Mann-Whitney *U* test. Differences in log_10_ CFU on catheters were assessed via a one-way ANOVA with Dunnett’s post-test (\**p* ≤ 0.05, \*\**p* ≤ 0.01, \*\*\*\**p* ≤ 0.0001).

***efaA* and *mntH2* are differentially transcribed during IE and CAUTI.** The results described above suggest that EfaCBA is the chief Mn transporter in bloodstream infections while MntH2 appears to play a more prominent role in CAUTI. Given the functional redundancy of these Mn transporters, we wondered if *efaCBA* and *mntH2* were differentially expressed when *E. faecalis* is in the bloodstream or in urine, thereby providing an explanation for the different phenotypic behaviors of Δ*efa* and Δ*mntH2* strains in different sites of infection. To address this possibility, we used quantitative reverse transcription-PCR (qPCR) to determine *efaA, mntH1* and *mntH2* transcription levels *in vivo.* When compared to cells grown in trypsinized beef heart medium, we found that *efaA* transcription was strongly induced in cells isolated from rabbit heart valves (~ 15-fold after two days and ~ 100-fold after three-four days) (Fig 10A). On the other hand, transcription of *mntH1* was moderately repressed after infection (7-fold and 3-fold repression after two and three-four days, respectively), whereas *mntH2* transcription remained largely unaltered over time (Fig 10A). Strikingly, *mntH2* was the only gene up-regulated (~ 10fold induction) in cells recovered from the bladders of mice 24 hours post-infection, while *efaA* and *mntH1* transcription was not significantly altered (Fig 10B). Since Mn plays a crucial role in promoting oxidative stress tolerance in lactic acid bacteria [6], in part by acting as the co-factor of superoxide dismutase (*sodA*), we also determined the transcription levels of *sodA* as a readout for oxidative stress in IE and CAUTI. Surprisingly, *sodA* transcription was downregulated (~ 15fold) at the early stage of endocarditis infection (two days) and did not differ from the inoculum condition at the later stage of infection (three-four days) (Fig 10A). Similarly unexpected, cells recovered from the bladders of infected mice displayed a ~ 25-fold reduction in *sodA* one day post-infection (fig 10B).

**Fig 10.**
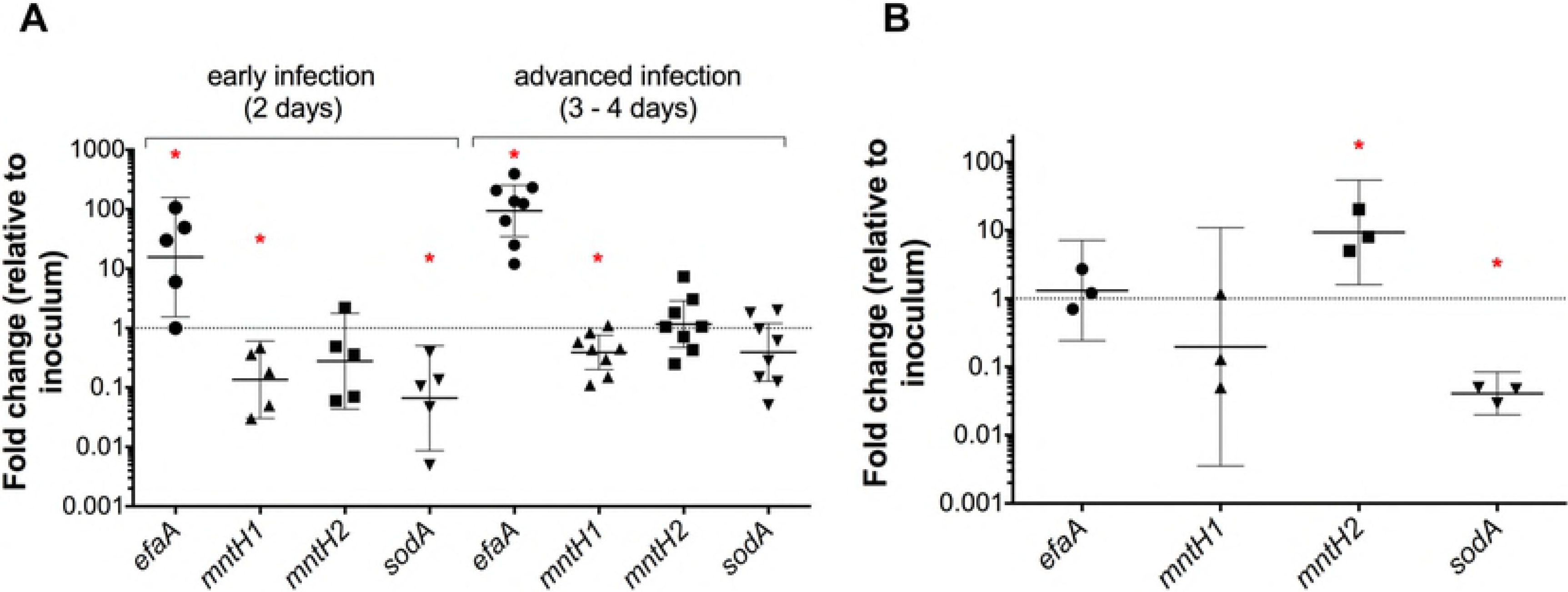
Transcription of *efaA, mntH1, mntH2* and *sodA* is differentially regulated during infection. OG1RF wild-type cells were harvested from (A) rabbit heart valves at 2 (early infection) and 3-4 (advanced infection) days post-infection (IE rabbit model, n = 5-8 independent animal samples) or from (B) mouse bladders 1 day post-infection (CAUTI mouse model, 3 pooled bladders/sample, n = 3). RNA was extracted, and transcript levels of *efaA, mntH1, mntH2*, and *sodA* were determined by qRT-PCR. The data points show fold-change of each individual sample relative to inoculum growth conditions. Horizontal lines and error bars denote geometric means and 95% confidence intervals. Confidence intervals that do not cross y = 1 (representing no fold-change in expression) are significantly different from the inoculum control (* *P*<0.05).

## Discussion

In this study we confirm previous *in silico* predictions that the *E. faecalis* core genome encodes three bona fide Mn transporters: one ABC-type (EfaCBA) and two Nramp-type transporters (MntH1 and MntH2) [27]. Early studies showed that the virulence of an *E. faecalis efaA* mutant was slightly delayed in a mouse peritonitis model despite the fact that the strain failed to display noticeable phenotypes *in vitro* [33, 36]. Here, we provide an explanation for such findings by showing that only the simultaneous inactivation of two or all three transporters can drastically impair Mn homeostasis under laboratory as well as *in vivo* conditions. Specifically, the Δ*efa*Δ*mntH1*Δ*mntH2* triple mutant strain was unable to grow (or grew very poorly) in Mn-restricted environments – a condition commonly encountered in human tissues – and was unable to robustly infect vertebrate and invertebrate animal hosts. To our knowledge, this is the first description of an *E. faecalis* mutant strain being virtually avirulent in multiple animal infection models. Moreover, this is also the first demonstration that the ability to acquire Mn using high affinity transporters within the urinary tract environment was shown to be an important aspect for the development of bacterial UTI.

While EfaCBA, MntH1 and MntH2 appear to be functionally redundant, simultaneous inactivation of *efaCBA* and *mntH2* often phenocopied the triple mutant strain. Specifically, the double Δ*efa*Δ*mntH2* mutant strain was highly sensitive to calprotectin, displayed biofilm formation defect in human urine and was virtually avirulent in the IE and CAUTI models. None of these phenotypes were further exarcebated in the triple mutant strain that also lacked *mntHl.* Considering that *mntHl* transcription is not as strongly affected by *in vitro* Mn fluctuations as *efaCBA* and *mntH2* [6, 34], it is tempting to speculate that EfaCBA and MntH2 are the primary high-affinity Mn transporters of *E. faecalis* responsible for maintaining Mn homeostasis in Mn-restricted environments. In contrast, MntH1 may serve as a housekeeping transporter, possibly with lower affinity for Mn than EfaCBA and MntH2, under Mn-replete conditions. The specificity and affinity (high/low) of each transport system for Mn, as well as other important transition metals such as Cu, Fe and Zn, will be determined in future studies.

Lactic acid bacteria like enterococci and streptococci are notoriously Mn-centric organisms, having a much higher nutritional demand for Mn than other bacterial groups [38]. This metabolic particularity might explain why the striking loss of virulence observed here contrasts with the moderate virulence attenuation of Mn transport mutants in gram-negative and other gram-positive pathogens [7, 8, 17]. Indeed, the complete loss of virulence of Δ*efa*Δ*mntH2* and Δ*efa*Δ*mntH1*Δ*mntH2* strains shown here is only comparable to the virulence defects of streptococcal strains lacking EfaCBA orthologs [25, 50, 51]. Interestingly, the observation that *E. faecalis* produces three highly conserved Mn transporters instead of two, such as *S. aureus* [17] and most streptococci [52], may be an indication that this opportunistic pathogen is better equipped to acquire Mn from the host environment than some of its closely-related pathogenic species. Alternatively, it is possible that *E. faecalis* may simply have a higher cellular demand for Mn than staphylococci and streptococci, having to rely on multiple Mn transporters to meet such high demand. This would resemble *Lactobacillus plantarum*, a non-pathogenic organism with five annotated Mn transporters (one P-type, one ABC-type, and three Nramp-type transporters) that has one of the highest cellular requirements for Mn among gram-positive and gram-negative bacteria [38, 53].

While Mn is the co-factor of several growth-promoting bacterial enzymes, Mn is thought to mediate bacterial virulence mainly by protecting cells from host-derived reactive oxygen species (ROS) [7, 8, 18, 43, 52, 54]. Previous studies have shown that Mn contributes to elimination of damaging ROS via three distinct mechanisms: (i) by direct non-enzymatic scavenging of superoxide radicals, (ii) by serving as the co-factor of the Mn-dependent superoxide dismutase (SOD) enzyme, and (iii) by replacing Fe as an enzymatic co-factor thereby protecting Fe-binding proteins from Fenton chemistry damage [8, 38, 55]. Not surprisingly, Mn transport mutants of different bacterial species display enhanced sensitivity to oxidative stresses *in vitro* and reduced macrophage survival [18, 19, 52, 56]. Global transcriptional analyses of *E. faecalis* grown in whole blood or urine *ex vivo* or isolated from a murine peritonitis model showed that transcription of several genes associated with ROS detoxification, including *sodA*, is induced when compared to cells grown in laboratory medium, indicating that bacterial cells have to cope with ROS stress during invasive infections [29–31]. Unexpectedly, we found that transcription of *sodA* was downregulated in the early stages of IE and CAUTI infection (Fig 10). While further studies are needed to determine if *E. faecalis* does not encounter oxidative stress during these infections, it is possible that the downshift in *sodA* is a response mechanism to lower the cellular Mn requirement when this nutrient is already restricted. In support of this possibility, *sodA* transcription has been previously shown to be repressed in *E. faecalis* during Mn limitation *in vitro* and in cells recovered from a rabbit subdermal abscess model 8 hours post-infection [6, 57]. Nevertheless, the discrepant transcriptional induction of *sodA* in different studies remains to be resolved. One possibility is that, during infection, oxidative stress fluctuates according to the dynamic and temporal immune responses mounted by the host.

By obtaining the transcriptional profile of *efaA, mntH1* and *mntH2* during IE and CAUTI, we found that the transcriptional responses of each system to different environmental cues *in vivo* can greatly differ. This was particularly noticeable for *efaA* and *mntH2*, since *efaA* was strongly induced in cells recovered from heart valves whereas *mntH2* was induced in CAUTI (Fig 10). In contrast, transcription of *mntH1* was repressed in heart valves while remaining largely unaltered in CAUTI. Considering that *efaCBA, mntH1* and *mntH2* have been previously shown to be regulated by EfaR [33], a metalloregulator from the DtxR family, the different transcriptional profiles of these genes were somewhat unexpected. However, the same study proposed that, similar to *Baciluus subtilis* and *S. enterica*, there could be other factors regulating transcription of Mn transport genes in *E. faecalis* [8, 33, 58, 59]. In fact, *in silico* analysis of the *E. faecalis* OG1RF genome identified two Fur-binding consensus sequences upstream *efaC*, the first gene in the *efaCBA* operon, indicating that this dual Fe/Mn transporter may also be regulated in response to Fe availability, oxidative stress or both. Future studies are warranted to identify the environmental cues present in blood and urine that trigger the different transcriptional responses of *efaCBA* and *mntH2*, as well as the potential *cis* and trans-acting elements regulating these responses.

In addition to differences in expression and, possibly, metal-binding affinity, the distinct behaviors of *efaCBA* and *mntH2* mutants in the context of IE and CAUTI may have additional explanations. For instance, EfaA homologs of *S. sanguinis* (SsaB), *S. parasanguinis* (FimA) and *S. pneumoniae* (PsaA) have been proposed to act as adhesins to a variety of relevant surfaces [52]. While the role of Mn ABC transport permeases as moonlighting proteins that participate in cell adhesion is a subject of debate [52], the possibility that EfaA can also function as a surface adhesin should not be completely excluded at this point. Alternatively, considering that the availability of free Mn and Fe is greatly restricted in the bloodstream, it is possible that the dual ability of EfaCBA to import both metals as suggested by cellular metal quantifications is physiologically relevant during bloodstream infections but not during UTI. In support of this possibility, both Fe and Mn are growth-limiting factors for *E. faecalis* in serum [6], while only Mn could fully rescue the growth defect of the triple Mn transport mutant in urine (Fig 7).

At this time, the dominant role of MntH2 in CAUTI is less clear. Recent work in *Streptococcus agalactiae* showed that the Nramp-type MntH is induced at low pH and facilitates macrophage and acid stress survival [60]. However, in the experiments reported herein, urine pH of infected mice remained stable at ~ 6.5 over the infection period (data not shown), suggesting that transcriptional induction of *mntH2* in urine cannot be attributed to a low pH response. Moreover, the Δ*mntH2* mutant was not more sensitive than the parent OG1RF strain when exposed to increasing concentrations of HCl in a disk diffusion assay (data not shown). Alternatively, we hypothesized that MntH2 might contribute to Cu detoxification during CAUTI. Specifically, Cu and ceruloplasmin - the major mammalian Cu-binding protein - were recently found to accumulate during bacterial UTI and uropathogenic *Escherichia coli* (UPEC) has been shown to up-regulate Cu efflux systems during clinical UTI to avoid Cu toxicity [61, 62]. Thus, we tested the ability of parent and Δ*mntH2* strains to grow in human urine supplemented with Cu. However, both strains were completely resistant to physiological concentrations of Cu (up to 0. 5 μm) and showed identical levels of Cu sensitivity at supra-physiological concentrations (Fig S3). While in need of further study, it appears that the prominent role of MntH2 in CAUTI could be simply attributed to its higher expression levels in the bladder environment when compared to EfaCBA.

In summary, here we show that maintenance of Mn homeostasis is an essential trait enabling *E. faecalis* to fully exert its full pathogenic potential. Specifically, simultaneous inactivation of *efaCBA* and *mntH2* renders *E. faecalis* avirulent in two mammalian infection models while only a triple Δ*efa*Δ*mntH1*Δ*mntH2* mutant strain was attenuated in the *G. mellonella* invertebrate model. These results expand the knowledge that bacterial Mn transport systems are promising targets for the development of novel antimicrobial therapies, which should be particularly effective to combat *E. faecalis* infections [63]. Work will soon be underway to uncover the significance of dietary Mn in the pathophysiology of *E. faecalis*, to elucidate the organ-dependent differential transcriptional regulation of each Mn transporter, and to understand the contribution of Mn-dependent responses to oxidative stress survival in the context of infection.

## Materials and Methods

**Bacterial strains, standard growth conditions and antibodies.** Bacterial strains and plasmids used in this study are listed in Table 1. All *E. faecalis* strains were routinely grown overnight at 37°C in BHI supplemented with 150 μM MnSO_4_. When required, 10 μg ml^−1^ erythromycin was added to the growth medium for stable maintenance of plasmids in the complemented strains. The primary antibodies used in the study were rabbit anti-*Streptococcus* group D antigen (anti-*E. faecalis* lipoteichoic acid) [64] and mouse anti-EbpA^Full^ [46]. Horseradish peroxidase (HRP)-conjugated goat anti-mouse and goat anti-rabbit antisera from KPL and IRDye 680LT goat anti-rabbit from LI-COR Biosciences were used as secondary antibodies.

**Table 1.** Bacterial strains and plasmids used in this study.

| Strains | Relevant characteristics | Source |
| --- | --- | --- |
| <u>*E. faecalis*</u> |  |  |
| OG1RF | Reference sequenced strain; Rif<sup>R</sup> Fus<sup>R</sup> | Laboratory stock |
| $\Delta efaCBA$ | *efaCBA* deletion | This study |
| $\Delta mntH1$ | *mntH1* deletion | This study |
| $\Delta mntH2$ | *mntH2* deletion | This study |
| $\Delta efaCBA \Delta mntH1$ | *efaCBA* deletion; *mntH1* deletion | This study |
| $\Delta efaCBA \Delta mntH2$ | *efaCBA* deletion; *mntH2* deletion | This study |
| $\Delta mntH1 \Delta mntH2$ | *mntH1* deletion; *mntH2* deletion | This study |
| $\Delta efaCBA \Delta mntH1 \Delta mntH2$ | *efaCBA* deletion; *mntH1* deletion;<br>*mntH2* deletion | This study |
| OG1RF + pTG001 | Empty vector, Erm<sup>R</sup> | This study |
| $\Delta efaCBA \Delta mntH1 \Delta mntH2$ + pTG001 | Triple mutant, empty vector, Erm<sup>R</sup> | This study |
| $\Delta efaCBA \Delta mntH1 \Delta mntH2$ + pTG-*efa* | Triple mutant, *efaCBA*<sup>+</sup>, Erm<sup>R</sup> | This study |
| $\Delta efaCBA \Delta mntH1 \Delta mntH2$ + pTG-*mntH1* | Triple mutant, *mntH1*<sup>+</sup>, Erm<sup>R</sup> | This study |
| $\Delta efaCBA \Delta mntH1 \Delta mntH2$ + pTG-*mntH2* | Triple mutant, *mntH2*<sup>+</sup>, Erm<sup>R</sup> | This study |
| CK111 | OG1Sp *upp4::P23repA4*, Spec<sup>R</sup> | [37] |
| <u>*E. coli*</u> |  |  |
| EC1000 | Host for RepA-dependent plasmids | [65] |
| DH10B | Cloning host for pTG001 | Laboratory stock |
| <u>Plasmids</u> |  |  |
| pCJK47 | Donor plasmid, carries P- *pheS* *; Erm<sup>R</sup> | [37] |

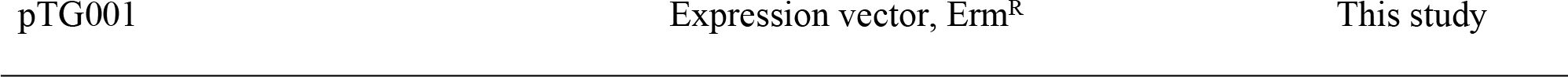

**Construction of Mn transport deletion strains.** Deletion of *efaCBA, mntH1,* and *mntH2* from the *E. faecalis* OG1RF strain was carried out using the pCJK47 markerless genetic exchange system [37]. Briefly, ~ 1 kb PCR products flanking the *efaCBA, mntH1*, or *mntH2* coding sequences were amplified with the primers listed in S1 Table. The amplicons included the first and last residues of the coding DNA to avoid unanticipated polar effects. Cloning of amplicons into the pCJK47 vector, electroporation and conjugation into *E. faecalis* strains and final isolation of single mutant strains were carried out as previously described [37]. Double mutants were obtained by conjugating the different pCJK constructs accordingly into Δ*efa*, Δ*mntH1* or Δ*mntH2* single mutants. A triple mutant was obtained by conjugating the pCJK-*mntH2* plasmid into the Δ*efa*Δ*mntH1* double mutant. All gene deletions were confirmed by PCR sequencing of the insertion site and flanking sequences in single, double and triple mutants. BHI agar was supplemented with 150 μM MnSO_4_ to enable isolation of the triple mutant strain.

**Construction of complementation strains.** The shuttle vector pTG001, a modified version of the nisin-inducible pMSP3535 plasmid [66] with an optimized RBS and additional restriction cloning sites (a gift from Anthony Gaca, Harvard Medical School), was used to complement the Δ*efa*Δ*mntH1*Δ*mntH2* triple mutant strain. Briefly, the coding sequence of *efaCBA, mntH1*, or *mntH2* was amplified from OG1RF (*efaCBA*, *mntH1*, *mntH2*) using the primers listed in S1Table, digested with the appropriate restriction enzymes and ligated into pTG001 digested with compatible restriction enzymes to yield plasmids pTG-*efa*, pTG-*mntH1* and pTG-*mntH2*. Upon propagation in *E. coli* DH10B, pTG001 (empty plasmid) and the complementation vectors were electroporated into OG1RF wild-type or Δ*efa*Δ*mntH1*Δ*mntH2* strains using a standard protocol [67] modified such that electroporated cells were immediately recovered in BHI supplemented with 0.4 M sorbitol. Presence of plasmids were confirmed via PCR amplification of the DNA insert region using plasmid-specific primers.

**Growth assays in Mn-depleted media.** *In vitro* growth of strains in Mn-depleted medium was performed using the chemically-defined FMC medium [68] depleted for Mn as previously described [6]. Overnight cultures were diluted 1:40 in complete FMC (metal-replete) and grown aerobically at 37°C to an OD_600_ of 0.25 (early exponential phase). Then, cultures were diluted 1:100 into fresh FMC depleted for Mn, and cell growth was monitored using the Bioscreen growth reader monitor (Oy Growth Curves AB Ltd.). To assess colony formation on BHI plates, overnight cultures were washed once in sterile PBS containg 0.1 mM EDTA to chelate extracellular divalent cations, followed by a second PBS wash to remove EDTA. Washed cells were diluted 1:100 in PBS, and 5 μl aliquots were spotted on plain BHI agar plates or BHI plates containing 150 μM MnSO_4_. Plates were incubated overnight at 37°C before growth was recorded. Growth in the presence of purified calprotectin and the His (103-105) - Asn calprotectin variant (a gift from Walter Chazin and Eric Skaar, University of Vanderbilt) was adapted from previous reports [15, 69, 70]. Briefly, overnight cultures were diluted 1:50 in fresh BHI broth and incubated 1 hour at 37°C. Then, cultures were diluted 1:100 into calprotectin medium, consisting of 38% BHI and 62% CP buffer [20 mM Tris pH 7.5, 100 mM NaCl, 3 mM CaCl_2_, 5 mM β-mercaptoethanol]. Native calprotectin and its variant unable to bind Mn [15] were added at a sub-inhibitory concentration (120 μg ml^−1^), and growth at 37°C was recorded at 30 min intervals for up to 24 hours.

**ICP-OES.** Total metal concentration in BHI, human urine and within bacterial cells was determined as previously described by <u>i</u>nductively <u>c</u>oupled <u>p</u>lasma - <u>o</u>ptical <u>e</u>mission <u>s</u>pectrometry (ICP-OES) at the University of Florida Institute of Food and Agricultural Sciences Analytical Services Laboratories [6]. Briefly, for quantification of metals in liquids, 18 ml BHI or urine were digested with 2 ml trace-metal grade 35% HNO_3_ prior to analysis. For quantification of the cellular metal content of bacterial cells, overnight cultures were washed twice with BHI and subsequently used to inoculate fresh BHI (1:20) and incubated statically at 37°C. After reaching an OD_600_ ~ 0.5, cells were harvested by centrifugation, washed twice in ice-cold PBS supplemented with 0.5 mM EDTA to chelate extracellular divalent cations, and aliquots were collected for metal analysis. Bacterial pellets were resuspended in 1 ml 35% HNO_3_ and digested at 90°C for 1 hour in a high-density polyethylene scintillation vial (Fisher Scientific). Digested bacteria were diluted 1:10 in reagent-grade H_2_O prior to ICP-OES metal analysis. Metal composition was quantified using a 5300DV ICP Atomic Emission Spectrometer (Perkin Elmer), and concentrations were determined by comparison with a standard curve. Metal concentrations were normalized to total protein content determined by the bicinchoninic acid (BCA) assay.

***G. mellonella* model of systemic infection.** Larvae of *G. mellonella* was used to assess virulence of OG1RF and its derivatives as described elsewhere [71]. Briefly, overnight cultures were washed to reduce Mn carryover. Groups of 15 larvae (200 - 300 mg in weight) were injected with 5 |il of bacterial inoculum containing ~ 5×10^5^ CFU. Larvae injected with heat-inactivated *E. faecalis* OG1RF (30 min at 100°C) were used as negative control. After injection, larvae were kept at 37°C and *G. mellonella* survival was recorded at selected intervals for up to 72 hours. Experiments were performed independently at least six times with similar results.

**Growth and survival in serum and urine.** Growth and survival of strains in pooled human serum (Lee Biosolutions) was monitored as previously described [6]. Briefly, overnight cultures grown in BHI+Mn were washed in sterile PBS as indicated above to remove excess Mn and inoculated 1:200 in human serum. Cultures were incubated aerobically at 37°C and, at selected time points, aliquots were serially diluted and plated on Mn-supplemented BHI plates for colony-forming unit (CFU) determination. For determination of growth in human urine, overnight cultures were normalized to an OD_600_ of 1.0 in BHI+Mn. Cells were washed 3 times with 1X PBS followed by 1:1000 dilution into undiluted pooled female urine supplemented with 20 mg ml^−1^ of BSA and incubated at 37°C. Bacterial growth was monitored by quantifying CFU as described above. When indicated, serum and urine were supplemented with a final concentration of 1 mM FeSO_4_ (99%, Sigma), 1 mM MnSO_4_ (99%, Sigma) or with increasing concentrations (0.5 μM, 0.5 mM or 1 mM) of CuSO_4_ (99%, Acros Organics). Urine was pooled from 3 healthy female donors, clarified by centrifugation, filter-sterilized, and adjusted to pH 6.5 prior to use. All urine samples were collected after obtaining written consent as per the study approval from the Washington University School of Medicine Internal Review Board (approval ID #201207143).

**Experimental rabbit IE model.** Male and female specific pathogen-free New Zealand White rabbits (2–4 kg; RSI Biotechnology) were utilized in an endocarditis model to assess virulence as previously described [25]. Prior to surgery, the rabbits were sedated and anesthetized with a cocktail of ketamine, xylazine and buprenorphine. A 19-gauge catheter was inserted into the aortic valve by way of the right carotid artery to induce minor damage. Each catheter was trimmed and sutured in place and the incision was closed with staples. Later the same day, duplicate inoculum preparation began by inoculating all strains into BHI for overnight growth in 6% O_2_. For the Δ*efa*Δ*mntH1*Δ*mntH2* strain only, BHI medium was supplemented with 100 μM Mn. The next morning, cultures were diluted 1:10 in fresh BHI with Mn supplementation and incubated with lids tight at 37°C until an OD_600_ of ~0.8 was obtained. The cells were then washed twice with 7 ml Chelex-treated (Bio-Rad) PBS containing 0.1mM EDTA. As determined in previous pilot experiments, a portion of the duplicate resuspended cells was removed and diluted to achieve equal strain representation and a total inoculum of ~10^7^ CFU/ml. From each sample, 1 ml was removed, sonicated and plated for enumeration, while 0.5 ml were injected into the peripheral ear vein of each of three rabbits per inoculum. A total of six animals were used for each set of experiments. However, one animal per group had to be euthanized before the set time point and was not included in the final analysis. Forty-eight hours after inoculation, rabbits were euthanized via intravenous injection of Euthasol (Med-Pharmex Inc.). Catheter placement was verified upon necropsy. Harvested cardiac vegetations and spleens were placed into PBS, homogenized and plated on Mn-supplemented BHI agar plates. Resulting colonies were picked and patched onto a new BHI plate supplemented with Mn as appropriate. PCR screens were used to determine the frequency with which each strain was recovered from heart vegetations and spleens. Thus, 50 to 100 colonies per organ and animal were randomly chosen and screened by colony PCR (primers listed in table S1) for their *efaCBA* and *mntH2* status (either wild-type or deleted gene).

**Mouse catheter implantation and infection.** The mice used in this study were 6-week-old female wild-type C57BL/6Ncr mice purchased from Charles River Laboratories. Mice were subjected to transurethral implantation and inoculated as previously described [47]. Mice were anesthetized by inhalation of isoflurane and implanted with a 5-mm length platinum-cured silicone catheter. When indicated, mice were infected immediately following catheter implantation with 50 μl of ~2 × 10^7^ CFU of bacteria in PBS introduced into the bladder lumen by transurethral inoculation as previously described [47]. To harvest the catheters and organs, mice were euthanized 24 hours post-infection by cervical dislocation after anesthesia inhalation, and bladder, kidneys, spleen and heart were aseptically harvested. Subsequently, the silicone implant was retrieved from the bladder.

**Biofilm assays on fibrinogen-coated silicone catheters and 96-well polystyrene plates.** Silicone catheters (1 cm, Nalgene 50 silicone tubing, Brand Products) or 96-well polystyrene plates (Grenier CellSTAR) were coated overnight at 4°C with 100 μg ml^−1^ human fibrinogen free of plasminogen and von Willebrand Factor (Enzyme Research Laboratory). The next day, *E. faecalis* overnight cultures were diluted to an optical density (OD_600_) of 0.2 in BHI broth. The diluted cultures were centrifuged, washed three times with 1× PBS, and diluted 1:100 in urine supplemented with 20 mg ml^−1^ BSA. Bacterial cells were allowed to attach to the fibrinogen-coated silicone catheters or 96-well polystyrene plates for 24 hours at 37°C under static conditions. After bacterial incubation, catheters and microplates were washed with PBS to remove unbound bacteria. Half of the catheters were vortexed for 30 sec, sonicated for 5 min, and vortexed again for 30 sec to retrieve bacteria in the biofilm for CFU quantification. The other half of catheters were used to visualize biofilms. Briefly, catheters were fixed with formalin for 20 min and then washed three times with PBS. Catheters were blocked at 4°C with 5% dry skin milk PBS, followed by three washes with PBS-T. After the washes, plates were incubated for an hour at room temperature with rabbit anti-*Streptococcus* group D antigen antisera (1:500) in dilution buffer. Plates were washed with PBS-T, incubated with the Odyssey secondary antibody (goat anti-rabbit IRDye 680LT, diluted 1:10,000) for 45 min at room temperature and washed three times with PBS-T. As a final step, plates were scanned for infrared signal using the Odyssey Imaging System (LI-COR Biosciences). Fibrinogen coated-catheters were used as a control for any auto-fluorescence. For assessment of biofilm formation on fibrinogen-coated 96-well polystyrene plates, microplates were stained with 0.5% crystal violet for 10 min at room temperature. Excess dye was removed by rinsing with sterile water and then plates were allowed to dry at room temperature. Biofilms were resuspended with 200 μl of 33% acetic acid and the absorbance at 595 nm was measured on a microplate reader (Molecular Devices). Experiments were performed independently in triplicate per condition and per experiment.

**Expression of EbpA on the cell surface of Mn transporter mutants.** Surface expression of EbpA by *E. faecalis* OG1RF parent and Mn transporter mutants was determined by ELISA as previously described [72] Bacterial strains were grown for 18 hours in urine supplemented with 20 mg ml^−1^ of BSA. Then, cells were washed three times with PBS, normalized to an OD_600_ of 0.5, resuspended in 50 mM carbonate buffer (pH 9.6) containing 0.1% sodium azide and used to coat (100 μl aliquots) Immulon 4HBX microtiter plates overnight at 4°C. The next day, plates were washed three times with PBS-T (PBS containing 0.05% Tween 20) to remove unbound bacteria and blocked for 2 hours with 1.5% BSA–0.1% sodium azide-PBS (BB) followed by three washes in PBS-T. EbpA surface expression was detected using mouse anti-EbpA^Full^ antisera, which was diluted 1:100 in PBS dilution buffer (PBS with 0.05% Tween 20, 0.1% BSA, and 0.5% methyl α-d-mannopyronoside) before serial dilutions were performed. A 100-μl volume was added to the plate, and the reaction mixture was incubated for additional 2 hours. Subsequently, plates were washed three times with PBS-T, incubated for 1 hour with HRP-conjugated goat anti-rabbit antisera (1:2,000), and washed again three times with PBS-T. Detection was performed using a TMB substrate reagent set (BD). The reaction mixtures were incubated for 5 min to allow color to develop, then the reactions were stopped by the addition of 1.0 M sulfuric acid. The absorbance was determined at 450 nm. Titers were defined as the last dilution with an A450 of at least 0.2. As an additional control, rabbit anti-*Streptococcus* group D antiserum was used to verify that whole cells of all strains were bound to the microtiter plates at similar levels. EbpA expression titers were normalized against the bacterial titers at the same dilution.

**Measurement of *efaA, mntH1, mntH2* and *sodA* transcription during enterococcal IE and CAUTI.** The aortic valve homogenate RNAs used for these experiments were collected and analyzed in another study from rabbits infected with *E. faecalis* OG1RF in an experimental model of IE (Colomer-Winter C, Gaca AO, Chuang-Smith ON, Lemos JA, and Frank KL, manuscript in preparation). Gene expression was studied in RNA samples collected from animals infected for two days (early infection, n=5 rabbits) or three to four days (advanced infection, n=8 rabbits). For transcript quantifications during CAUTI, OG1RF cells were recovered from bladders of three infected mice, pooled and immediately placed in RNAlater solution (in triplicate, total n=9 animals). RNA extraction, reverse transcription and real-time PCR were carried out following standard protocols [6, 73].

**Statistical Analysis.** Data were analyzed using GraphPad Prism 6.0 software unless otherwise stated. Differences in cellular metal uptake, final growth in Mn-depleted FMC or human urine, and the percentage of strains recovered from the rabbit IE experimental model were determined via ordinary one-way ANOVA followed by post-test comparisons. Log_10_-transformed CFU values from serum survival and calprotectin growth experiments were analyzed via a two-way ANOVA followed by comparison post-tests. While differences in the *G. mellonella* killing rate of mutant strains was assessed with the Mantel-Cox log-rank test, final 72 hour survival of larvae were similarly compared via ordinary one-way ANOVA with Dunnett’s multiple comparison post-test. Of note, two outliers (one for a Δ*mntH1* replicate, another for a Δ*efa*Δ*mntH1*Δ*mntH2* replicate) were identified using the ROUT method (Q = 1%) and removed from the final analysis. For CAUTI experiments, biofilm assays and EbpA expression, data from multiple experiments were pooled and Two-tailed Mann-Whitney *U* tests were performed. To determine statistical significance in fold-change transcription of selected genes, fold-change values for each gene were plotted as geometric mean with the corresponding 95% confidence interval (error bars). A line at y = 1 denotes equal transcripts in inoculum control vs *in vivo* condition. 95% confidence intervals that did not cross y = 1 were significantly different (* p<0.05) from the inoculum control.

**Ethics Statement.** Urine samples were collected after obtaining informed written consent for all subjects enrolled in the study as per the study approval from the Washington University School of Medicine Internal Review Board (approval ID #201207143). Populations generally classified as vulnerable, including children under the age of 18, were not enrolled in the study. Only subjects 18 years of age or older at the time of consent were eligible for the study. No group or persons were excluded from the study due to race, ethnicity or gender. Case subjects were enrolled solely based on the eligibility criteria.

Animal procedures for rabbit IE were approved by the Institutional Animal Care and Use Committee of the Virginia Commonwealth University as part of protocol number AM10030 as well as the University of Minnesota Institutional Animal Care and Use Committee as part of protocol number 0910A73332. The Washington University Animal Studies Committee approved all mouse infections and procedures as part of protocol number 20150226. All animal care was consistent with the Guide for the Care and Use of Laboratory Animals from the National Research Council and the USDA Animal Care Resource Guide.

## Acknowledgments

The modified shuttle-vector pTG001 for construction of complementation strains was a gift from Dr. Anthony O. Gaca at Harvard Medical School. Wild-type calprotectin and calprotectin ΔMn Tail mutant [His (103-105)-Asn] were generously provided by Dr. Eric Skaar and Dr. Walter Chazin at Vanderbilt University. We thank Dr. Jessica Kajfasz for critical reading of the manuscript and Dr. Cara Olsen at the USUHS Biostatistics Consulting Center for valuable input on the statistical analysis of *in vivo* qPCR data.

## Supporting information

**S1 Fig. *G. mellonella* survival rates.** Kaplan-Meyer plots of *G. mellonella* larvae injected with *E. faecalis* OG1RF or the single, double, or triple mutants. Each Kaplan-Meyer plot is a representative of an experiment repeated at least six independent times. Differences in the *G. mellonella* killing rates of the mutant strains compared to the parent OG1RF strain were assessed with the Mantel-Cox log-rank test. (** *p* ≤ 0.01).

**S2 Fig. Systemic dissemination of *E. faecalis* strains during CAUTI.** OG1RF and its derivatives were inoculated into the bladder of mice immediately after catheter implantation. After 24 hours, animals were euthanized and bacterial burdens in (A) kidneys, (B) spleens and (C) hearts determined. Graphs show total CFU recovered from these sites, and each symbol represents an individual mouse. Symbols on the dashed line indicate that recovery was below the limit of detection (LOD, 40 CFU). Data were pooled from at least two independent experiments, and median value is shown as a horizontal line. Statistical differences were analyzed by a twotailed Mann-Whitney *U* test (\**p* ≤ 0.05, \*\**p* < 0.005, \*\*\**p* < 0.001).

**S3 Fig. MntH2 does not confer tolerance to Cu toxicity in urine.** Survival of OG1RF wild-type and Δ*mntH2* strains in human urine supplemented with (A) 0.5 μM, (B) 0.5 mM, or (C) 1 mM CuSO_4_. Control corresponds to urine without CuSO_4_ supplementation. Aliquots at selected time points were serially diluted and plated on BHI + Mn plates for CFU enumeration. Survival of strains was recorded over 48 hours. The graphs show the average log_10_-transformed CFU mean and standard deviations of at least three independent experiments Differences were assessed via two-way ANOVA with Tukey’s post-test (\*\**p* ≤ 0.01, \*\*\**p* ≤ 0.001, \*\*\*\**p* ≤ 0.0001).

**S1 Table.** Primers used in this study.

